# A Meta-analysis of the known Global Distribution and Host Range of the *Ralstonia solanacearum* Species Complex

**DOI:** 10.1101/2020.07.13.189936

**Authors:** Tiffany M. Lowe-Power, Vienna Elmgreen, Jason Avalos, Yali Bai, Maria Charco Munoz, Kyle Chipman, Willem Elings, Larissa Hayes, Miranda Jaramillo, Neha Prasad, Benjamin Ramirez, Ajay Sandhar, Cloe E. Tom, Darrielle Williams

## Abstract

The *Ralstonia solanacearum* species complex is a group of genetically diverse plant wilt pathogens. We created a database that documents the reported global distribution and host range of *Ralstonia* clades (e.g. phylotypes and sequevars). In this sixth release, we have cataloged information from 353 sources that report one or more *Ralstonia* strains isolated from 113 countries or territories. Metadata for nearly 14,000 strains are available as a supplemental table. We continue to maintain this database and the RSSC KBase Narrative. More detailed information regarding specific methods can be found in previous versions of this living pre-print.

Recent reports expand the known distribution of several lineages: *Ralstonia syzygii* subsp. *syzygii* in Brazil, *Ralstonia syzygii* subsp. *celebesensis* (Banana Blood Disease lineage) in continental Asia, and phylotype I in Europe.

The interactive *Ralstonia* Diversity Dashboard https://ralstoniadashboard.shinyapps.io/RalstoniaWiltDashboard/) has been updated with this new version of the dataset. Additionally, this version of the living pre-print has new visualizations.

## Introduction

Bacterial pathogens in the *Ralstonia* species complex are xylem pathogens that infect a broad range of agricultural and natural plant hosts. *Ralstonia* strains clog plant xylem vessels, leading plant hosts to wilt [1]. Historically, *Ralstonia* strains were classified based on carbon utilization patterns (“Biovar”) and host range (“Race”). However, DNA sequence-based taxonomies more accurately reflect the evolutionary trajectories of *Ralstonia* lineages.

Currently, the *Ralstonia solanacearum* species complex (RSSC) is classified into three species: *R. solanacearum*, *R. pseudosolanacearum*, and *R. syzygii*. The separation of *Ralstonia* into three species was first proposed by Remenant et al. 2010 [2], formalized by Safni et al. 2014 [3], and reinforced by Prior et al. 2016 [4]. Strains are also classified into a phylotype system, which overlaps with the species boundaries. All *R. solanacearum* strains are within phylotype II, but phylotype II is divided into IIA, IIB, and IIC groups. *R. pseudosolanacearum* strains are either in phylotype I or phylotype III. *R. syzygii* strains are in phylotype IV. Strains are further sub-classified into sequence variants, or “sequevars”, based on the DNA sequence of the conserved *egl* endoglucanase gene (**Fig. 1**). Because *Ralstonia* are known to be naturally competent, we hypothesized that horizontal gene transfer of *egl* could confound sequevar-based phylogenies. We recently used whole genome phylogenetic trees to interrogate the robustness of *egl*-based trees [5]. We found that *egl* trees work well for phylotype II, but *egl* trees are highly inaccurate for phylotype I. Although only a small number of phylotype III and IV genomes were analyzed, the longer branch lengths in phylotype III and IV suggest that *egl*-based trees estimate these strains’ phylogenetic position effectively. Since the phylotype-sequevar system was first developed and described by Prior and Fegan in 2005 [6], hundreds of papers have used this system to describe the genetic diversity of *Ralstonia* isolates around the world. However, no public database previously aggregated this population genetics information. Here we report the sixth release of such a database, and we use it to describe the host range and global distribution of each *Ralstonia* clade.

**Fig 1.**
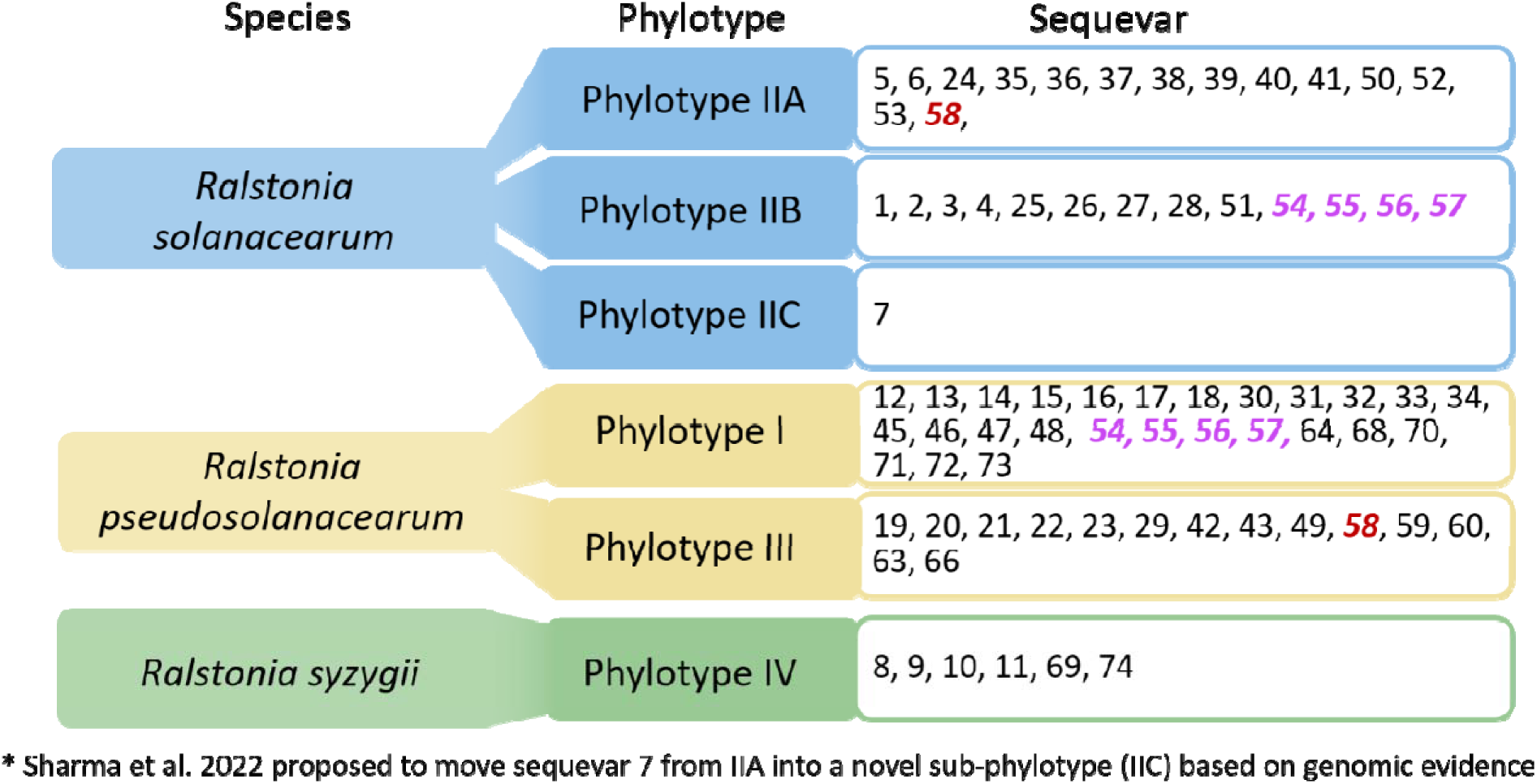
Taxonomic classification of the *Ralstonia* species complex at the species, phylotype and sequevar levels. Sequevars highlighted in red or purple italics are assigned to strains in conflicting phylotypes. As a note, many of the phylotype I sequevars are polyphyletic (See Sharma et al. 2022 for details [5]), so the *Ralstonia* community needs to identify a cost-effective alternative for describing diversity of these strains. The clonal lineages that cause Blood Disease of Banana (BDB) are nested within phyl. IV-10 with non-BDB phyl. IV strains.

### Updates from v5 to v6 of the *Ralstonia* diversity dataset

This section summarizes the changes between versions 5 and 6 of the “*Ralstonia* Diversity Dataset,” the meta-analysis we began in 2020 to document the known host range and global distribution of each sequevar in the *Ralstonia* species complex. In 2025, we built the *Ralstonia* Wilt Dashboard as an accompanying visualization and exploration tool to the database. The Dashboard is described in a publication [7] and is available at https://ralstoniadashboard.shinyapps.io/RalstoniaWiltDashboard/ and the short URL mirror: https://tinyurl.com/ralsto-dashboard. We intend to update this preprint at regular intervals as we populate the underlying *Ralstonia* Diversity Dataset.

In version 6 of the Dataset (this update), we incorporate data from 49 new sources and we maintain the user-friendly KBase narrative [8] established in version 4. Changes are summarized in **Fig 2** and **Table 1**. Th dataset is available from this *bioRxiv* preprint.

**Fig 2.**
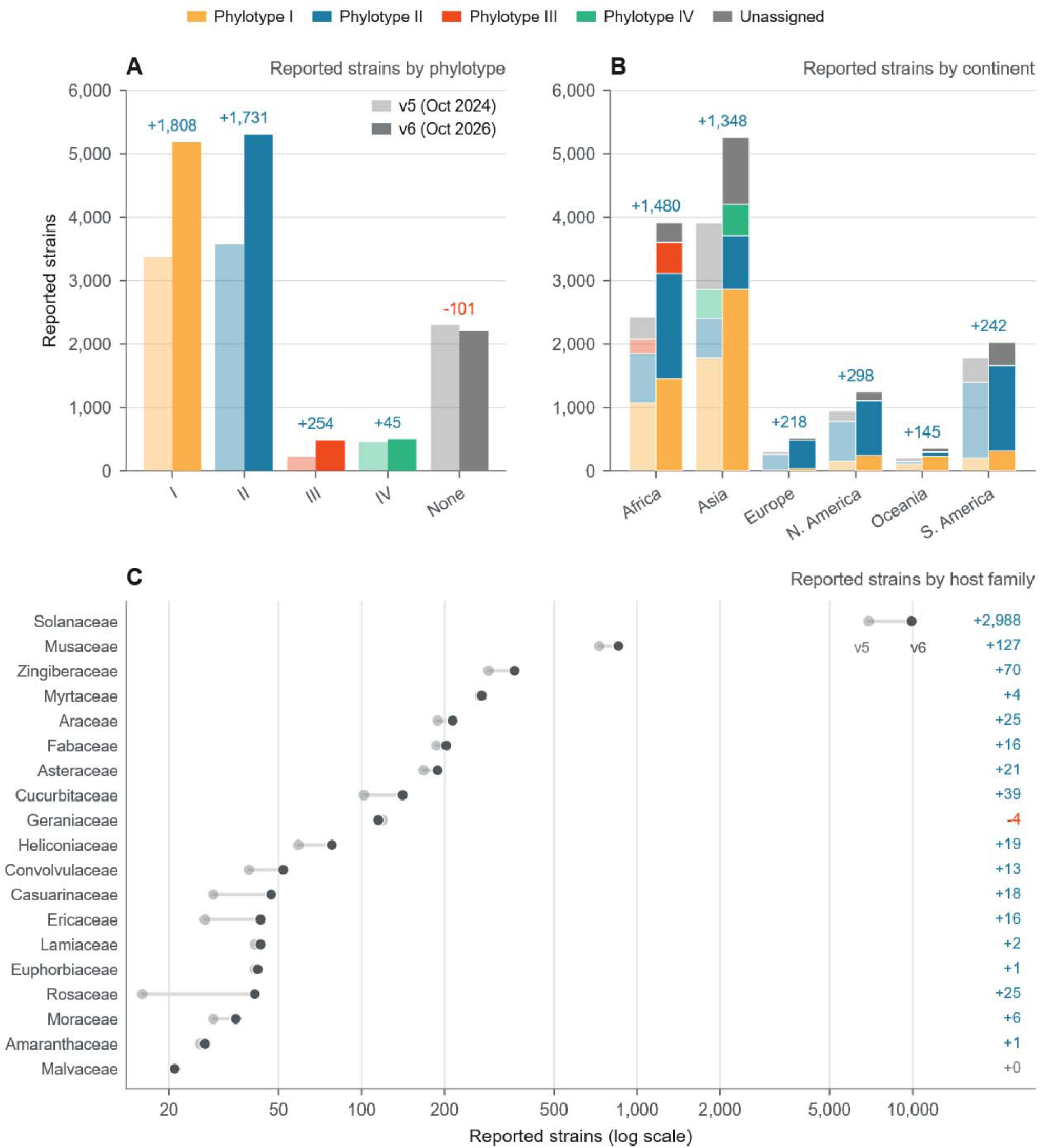
Summary of Database Changes in Version 6. Number of reported strains by (A) phylotype, (B) continent, and (C) plant families that correspond to 20 or more records in version 6. The light colors show values from version 5 (released 2024) and bold colors show version 6 (released 2026).

**Table 1:** Summary of the *Ralstonia* diversity dataset.

| Version | Database Release Date | # Sources<sup>a</sup> | Paper published range | # Strains | # Sequevars<sup>b</sup> | # Countries or Territories | # Host plants |
| --- | --- | --- | --- | --- | --- | --- | --- |
| v1 | 2020/07/03 | 35 | 2017-2020 | 1,625 | 57 | 50 | 47 |
| v2 | 2021/03/23 | 73 | 2012-2020 | 3,395 | over 64 | 86 | 94 |
| v3 | 2021/11/03 | 93 | 2007-2020 | 4,924 | Over 71 | 86 | 122 |
| v4 | 2022/08/30 | 197 | 1971-2022 | 7,794 | 68 | 105 | 414 |
| v5 | 2024/10/01 | 304 | 1971-2024 | 9,931 | 68 | 107 | 448 |
| v6 | 2026/10/31 | 353 | 1971-2026 | 13,668 | 69 | 113<sup>c</sup> | 465 |

<sup>a</sup> Most sources are publications, but others are databases or strain collections. Thus, the number of sources exceeds the number of papers in the List of Papers table.
<sup>b</sup> Note that sequevar is an imperfect phylogenetic system for phylotype I. See Sharma et al. 2022 for details [5]. An international collaboration has been initiated to create a genome-based naming system for sequevar-scale genetic diversity in RSSC.
<sup>c</sup> The database now includes reports of RSSC in Cambodia, Argentina, Russia, and Switzerland.

We added an “alternative strain names” column to this version of the database, which helped us consolidate duplicate entries for isolates archived in multiple collections under different strain identifiers. A future version will consolidate these entries more systematically. We are working with others in the *Ralstonia* community to cross-reference strain lists and create a master list of alternative names for strains.

## Methods

### Article Selection Criteria and Search Strategy

We prioritized articles that used the phylotype and/or phylotype-sequevar system to characterize strains. We identified these articles by using Google Scholar to find the papers and theses that cite “How complex is the *Ralstonia solanacearum* species complex?” by Fegan and Prior 2005 [6]. We did not include the Race or Biovar information in the database. Although host range is an important trait of strains, measuring multi-host virulence is labor and time-intensive, and has rarely been performed. Therefore, “race” assignments are usually listed solely based on the host-of-isolation, which does not accurately predict the strains’ host range. We also excluded biovar data because the carbon utilization patterns are not predictive of phylogeny or the most important phenotypes for *Ralstonia*. Further critique of these outdated classification schemes is detailed in [5]. To better estimate the diversity of potential hosts for *Ralstonia*, we searched the literature using terms such as “weeds”, “host range”, and older names for the pathogen (*Pseudomonas solanacearum*, *Burkholderia solanacearum*).

**Table 2:** List of Papers in this Study.

| Year Published | References |
| --- | --- |
| 1971 | [9,10] |
| 1976 | [11] |
| 1978 | [12] |
| 1980 | [13] |
| 1984 | [14] |
| 1986 | [15–25] |
| 1993 | [26–38] |
| 1995 | [39] |
| 1998 | [40–51] |
| 1999 | [52–54] |
| 2000 | [55] |
| 2001 | [56–59] |
| 2002 | [60,61] |
| 2003 | [62–64] |
| 2004 | [65,66] |
| 2005 | [67–70] |
| 2006 | [71–74] |
| 2007 | [75–83] |
| 2008 | [84–90] |
| 2009 | [91–100] |
| 2010 | [101–108] |
| 2011 | [109–124] |
| 2012 | [124–139] |
| 2013 | [140–155] |
| 2014 | [3,156–171] |
| 2015 | [172–188] |
| 2016 | [4,189–202] |
| 2017 | [203–220] |
| 2018 | [221–236] |
| 2019 | [237–248] |
| 2020 | [97,249–271] |
| 2021 | [272–285] |
| 2022 | [286–295] |
| 2023 | [296–311] |
| 2024 | [312–331] |
| 2025 | [332–350] |
| 2026 | [351–353] |
| Pre-prints | [354,355] |

### Converting strain metadata into a structured format

We focused on cataloging information that is relevant to the epidemiology of the *Ralstonia* strains: Phylogenetic position, host of isolation, and geographic location where isolated. Additionally, when listed, we include any NCBI accessions for the genome, *egl* marker genes, or other genes.

For **phylogeny**, we record the phylotype (I-IV), sub-phylotype (for phylotype II, this is the IIA, IIB, or IIC subdivision), and sequevar (1-74). Several sequevars have been subdivided based on phylogeny and/or ecologically important traits, so we created a “sub-sequevar” column to denote these. This includes multiple subdivisions of sequevar IIB-4 and IV-10 sequevar which includes the “*R. syzygii* subspecies *celebesensis*” (causes Blood disease of banana) and a clade of the paraphyletic “*R. syzygii* subspecies *indonesiensis*”.

For **host**, we created several columns to annotate the host at multiple taxonomic scales (order to species). We attempted to systematize all common names and species into a unified label. For example, all potato isolates are listed as *“Solanum tuberosum* (Potato)”. The systematic structure will make it easier to use R, Python pandas, or Excel equations to summarize and visualize the contents of the database.

For **location**, we record the most precise location information available (e.g. city, province, or country). We also have columns that describe the location at the country (or territory if the land region is geographically separated from the governing body), subcontinent, and continent levels.

### Curation, quality control, and correction of strain metadata

We curated records using Google Scholar and the NCBI Genome database. We refined both searches using keywords including *‘Ralstonia solanacearum’, ‘Ralstonia pseudosolanacearum’, ‘Ralstonia syzygii’, ‘Ralstonia pseudomonas’*, and *‘Ralstonia’.* We reviewed any paper that mentioned isolation of a strain within the *Ralstonia solanacearum* species complex and extracted metadata from it. At minimum, we collected ‘Host isolated from’ or ‘Location isolated’ from each paper. We additionally restricted the search using the ‘Sort by Year’ function. Using these methods, we reviewed papers and recorded metadata into individual Excel sheets. To avoid duplicated papers, we compared citations and did not record overlaps onto the master sheet. We then merged metadata from individual datasheets into the master metadata sheet.

We performed quality control and correction of strain metadata in OpenRefine and manually in Excel. OpenRefine is a free data management tool that applies changes across whole columns and datasets. OpenRefine includes many baseline functions that fix common entry mistakes by transforming data in bulk. Some of the most useful functions were ‘trim leading and trailing whitespaces’, ‘collapse consecutive whitespace’, and ‘cluster and edit’. The ‘Cluster and edit’ function groups together similar strings of characters such as ‘NCBI 5’ and ‘NCBI-5’. Using ‘cluster and edit’ paired with ‘facet’ functions, we grouped together all values that were qualitatively the same, such as *‘Solanum tuberosum’* and *‘Solanum tuberosum* (Potato)’ and ‘Potato’. Using this method, we consolidated values across all characterizations of the metadata. Manual corrections of strain metadata were made in Excel, namely host and location data. We expanded missing host taxonomic data from species name to genus, family, and order. We similarly expanded location data to country, subcontinent, and continent. We then geocoded location data using a custom R script with the MapBox API.

### Database

The full dataset is available as a supplementary table on this preprint.

### Extracting metadata for genomes available on NCBI

Genomes on NCBI do not always have a corresponding Genome Announcement or other publication, but most of these genomes are deposited with ecological metadata on the corresponding “BioSample” page. The NCBI taxonomy is disorganized because of taxonomic revisions in this group. Many genomes for *“Ralstonia pseudosolanacearum”* (taxonomy ID: 1310165) and *“Ralstonia syzygii”* (taxonomy ID: 28097) were deposited as the *“Ralstonia solanacearum”* (taxonomy ID: 305), primarily because they were deposited prior to the renaming in 2014. Moreover, several genomes of the blood disease pathogen *(Ralstonia syzygii* subsp. *celebesensis*) were deposited with their own taxonomy ID due to a peculiar taxonomic rule: these strains could not be formally named because the type strain was not viable in culture collections.

In July 2022, we extracted metadata for all RSSC genomes that we could identify in relevant “taxonomy IDs”. For each genome, we listed the “RefSeq Assembly Accession” when possible (GCF_xxx). Some assemblies have been excluded from RefSeq due to problems like “too many frameshifted proteins” or “highly fragmented genome”. In this case, we list the GenBank Assembly Accession (GCA_xxx). As of Summer 2024, many of those RefSeq genomes have been suppressed despite the genome being high quality and, sometimes, complete. Unfortunately, while conducting newly implemented quality analyses, the NCBI RefSeq staff suppressed the RefSeq genomes for all phylotype I, III, and IV strains that had been uploaded in the species *“Ralstonia solanacearum”* because these genomes were flagged as “unverified source organism”. As of summer 2024, NCBI has developed automated pipelines to check and report useful information on the new “NCBI Datasets” page: (1) assembly completeness/contamination with checkM and (2) taxonomic assignment based on ANI to type strains.

### Importing genomes into the “RSSC Phylogenomics” narrative on KBase

KBase is a user-friendly graphical user interface for biological informatics that is developed and maintained with significant investment from the US Department of Energy [8]. Because RSSC genomes are dispersed across several (often inaccurate) taxid pages, we developed a KBase Narrative that can help us and other scientists identify the phylogenetic identity of new RSSC genomes.

We used the structured data on our spreadsheet to assign a systematic name to each genome, such as: “IIC-7_UW853_UCD541_Tomato-Solanaceae_UnitedStates_GCA_036543205.1”. The systematic name lists:

1. Phylogenetic information (phylotype, sequevar, sub-sequevar)
2. Strain name (spaces and parentheses removed. Colons and hyphens were often removed, but underscores were often kept)
3. Host of isolation (common name and family)
4. Location (country/territory level)
5. Genome accession

Initially, there were genomes that lacked precise phylogenetic information. To identify the phylogenetic identity of these genomes, we used the KBase App “Batch Create GenomeSet v1.2.0” to create a GenomeSet object that included all genomes in the narrative. Then we used the “Insert Set of Genomes into SpeciesTree v2.2.0” to create a low-resolution phylogenetic tree. This app uses FastTree2 [356] with the - fastest setting to construct a tree based on an MSA of 49 universal bacterial genes. We inspected the tree, and we were able to assign many genomes to phylotypes and sequevars.

Wherever possible, we use the NCBI RefSeq or NCBI GenBank version of the genomes. However, when only the raw reads were available on SRA, we carried out a pipeline of importing the reads from SRA, assessing read quality with FastQC, trimming reads with Trimmomatic (Illumina) with settings to trim adapters when necessary or Filtlong (Nanopore when Illumina short reads were present), assessing trimmed reads with FastQC, assembling with SPAdes (Illumina) or Unicycler (Nanopore and Illumina), and annotation with Prokka.

### Data visualization

Within this pre-print, we used Claude to generate most of the visualizations. Claude used matplotlib 3.11 and Natural Earth 1:110m via the world-atlas package.

## Results and Discussion

We compiled 13,668 strains from 353 sources. The strains were isolated from 113 countries or territories (**Table 1**). Most of the reported strains are phylotype I or II strains (**Fig. 3**). We analyzed patterns in the host range and biogeographic distribution of the RSSC and of each phylotype. The full dataset is in **Table S1**.

**Fig 3.**
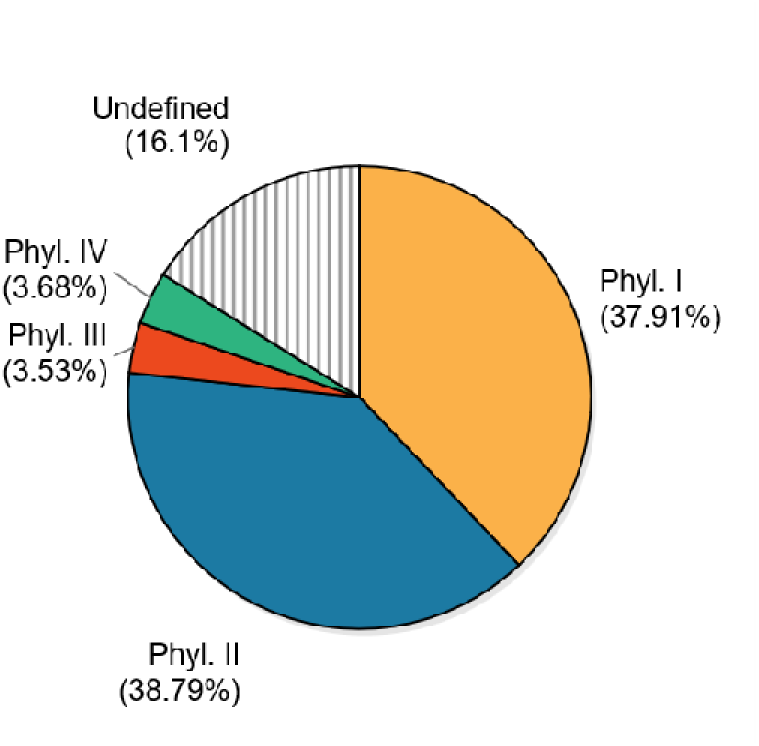
Phylotype assignments for strains in the Global *Ralstonia* Diversity database

### Host Range

Strain metadata varies in the specificity of the plant host. Some hosts are reported by only common name, others by species, others at just the genus level (e.g. *Musa* sp.) or family level (e.g. Solanaceae sp.). In earlier versions of the living pre-print, we counted *“Musa* sp.” as a distinct host from *“Musa acuminata”.* In this version, the tables and summaries count a “Genus sp.” as unique only if no species of that genus is listed elsewhere in the database. We applied the same logic to strains reported at the family level. We used Claude to consolidate these records. To enable comparisons of the different versions of the *Ralstonia* Diversity Dataset, we applied this approach retroactively to the counts listed in **Table 1**.

**Table 3:** The number of unique plant hosts (reported at various taxonomic levels)

|  | Species* | Genus | Family | Order | Countries |
| --- | --- | --- | --- | --- | --- |
| All RSSC | 465 | 280 | 85 | 33 | 113 |
| Phylotype I | 123 | 95 | 48 | 27 | 72 |
| Phylotype II | 93 | 60 | 36 | 25 | 81 |
| Phylotype III | 17 | 9 | 6 | 6 | 20 |
| Phylotype IV | 16 | 11 | 8 | 8 | 12 |
| Taxa only recorded for strains without phylotype assignments† | 277 | 153 | 29 | 3 | 7 |

\* The “Species” column includes some plants only identified to the genus level, e.g. “*Rosa* sp.”
† This row counts taxa, genera, families, orders, and countries recorded only for strains lacking phylotype assignments. Treat these with caution because it is possible the references for such hosts have mistakenly identified RSSC as the pathogen.

**Table 4:** The top 20 most common host species to be listed in the database, sorted by frequency.

| Host taxon | Strains | % of total | I | II | III | IV | None |
| --- | --- | --- | --- | --- | --- | --- | --- |
| *Solanum tuberosum* (Potato) | 4,200 | 32.19% | 431 | 2,984 | 299 | 151 | 335 |
| *Solanum lycopersicum* (Tomato) | 2,784 | 21.34% | 1,471 | 814 | 79 | 28 | 392 |
| *Solanum melongena* (Eggplant) | 834 | 6.39% | 677 | 48 | 15 | 0 | 94 |
| *Nicotiana tabacum* (Tobacco) | 810 | 6.21% | 710 | 34 | 6 | 1 | 59 |
| *Capsicum annuum* (Pepper) | 688 | 5.27% | 481 | 37 | 12 | 2 | 156 |
| *Musa acuminata* (Banana) | 494 | 3.79% | 3 | 316 | 0 | 77 | 98 |
| *Zingiber officinale* (Ginger) | 275 | 2.11% | 182 | 5 | 0 | 0 | 88 |
| *Musa acuminata* x *Musa balbisiana* (Blue Java) | 175 | 1.34% | 0 | 1 | 0 | 174 | 0 |
| *Eucalyptus pellita* (Large-fruited Red Mahogany) | 171 | 1.31% | 53 | 110 | 0 | 0 | 8 |
| *Solanum macrocarpon* (African Eggplant) | 138 | 1.06% | 132 | 0 | 5 | 0 | 1 |
| *Arachis hypogaea* (Peanut) | 99 | 0.76% | 60 | 4 | 0 | 0 | 35 |
| *Musa paradisiaca* (Plantain) | 81 | 0.62% | 1 | 70 | 0 | 1 | 9 |
| *Solanum dulcamara* (Bittersweet Nightshade) | 81 | 0.62% | 2 | 76 | 0 | 0 | 3 |
| *Solanum aethiopicum* (Bitter Tomato) | 68 | 0.52% | 62 | 6 | 0 | 0 | 0 |
| *Epipremnum aureum* (Pothos) | 52 | 0.40% | 0 | 17 | 0 | 0 | 35 |
| *Casuarina equisetifolia* (Ironwood Tree) | 47 | 0.36% | 40 | 3 | 0 | 0 | 4 |
| *Syzygium aromaticum* (Clove) | 46 | 0.35% | 1 | 2 | 0 | 40 | 3 |
| *Solanum nigrum* (Black Nightshade) | 44 | 0.34% | 13 | 17 | 2 | 0 | 12 |
| *Rosa* sp. (Rose) | 38 | 0.29% | 28 | 5 | 0 | 0 | 5 |
| *Solanum americanum* (American Black Nightshade) | 36 | 0.28% | 19 | 15 | 0 | 0 | 2 |

**Fig 4.**
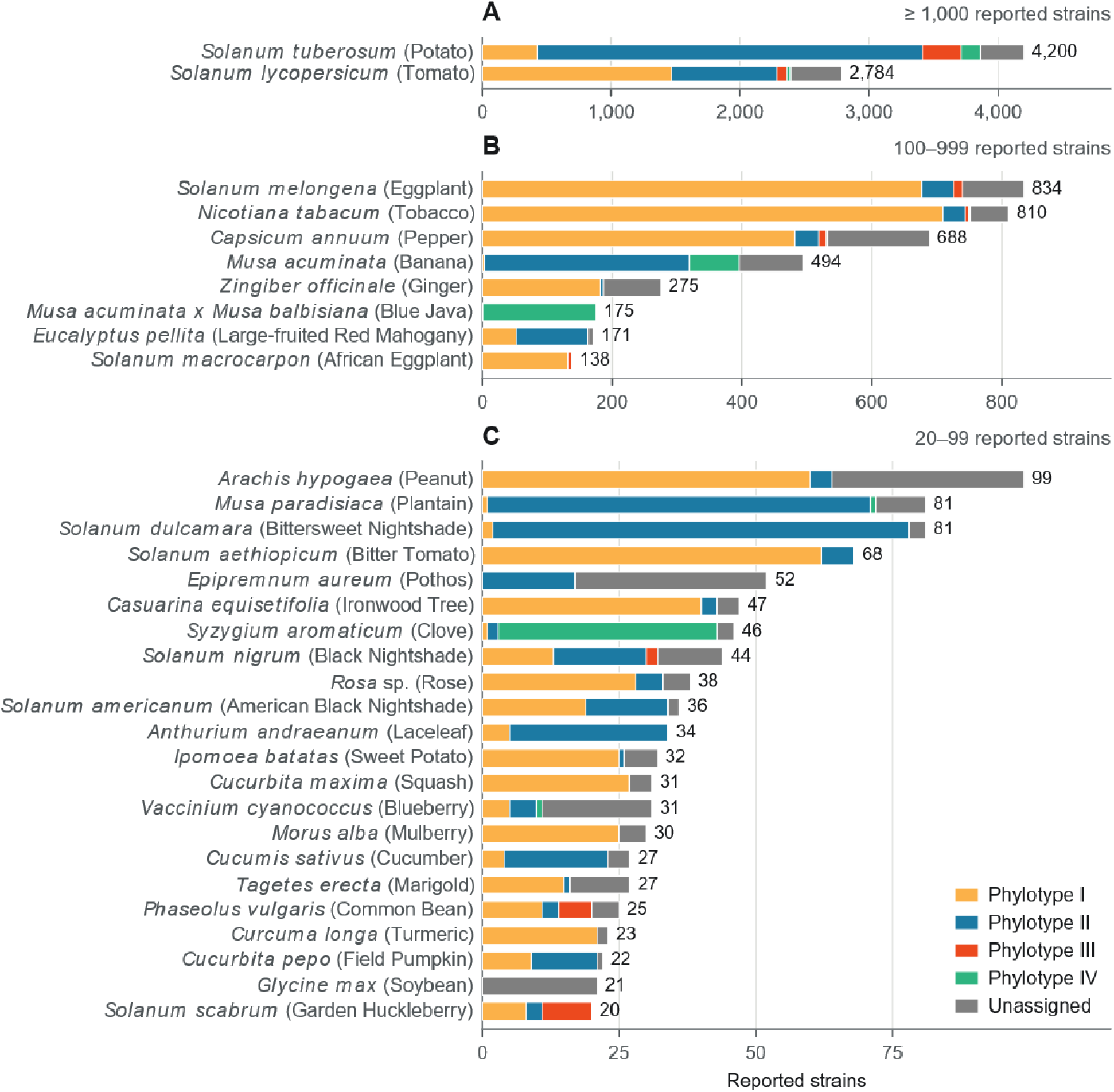
Phylotype composition of host species with 20 or more reported RSSC isolations. Bars give the number of reported strains of each phylotype, with the total printed at the right. Each panel has the axis scaled to a different maximum: (A) 1,000 or more strains, (B) 100– 999, (C) 20–99. Fig. 5 below is a companion to this figure.

**Fig 5.**
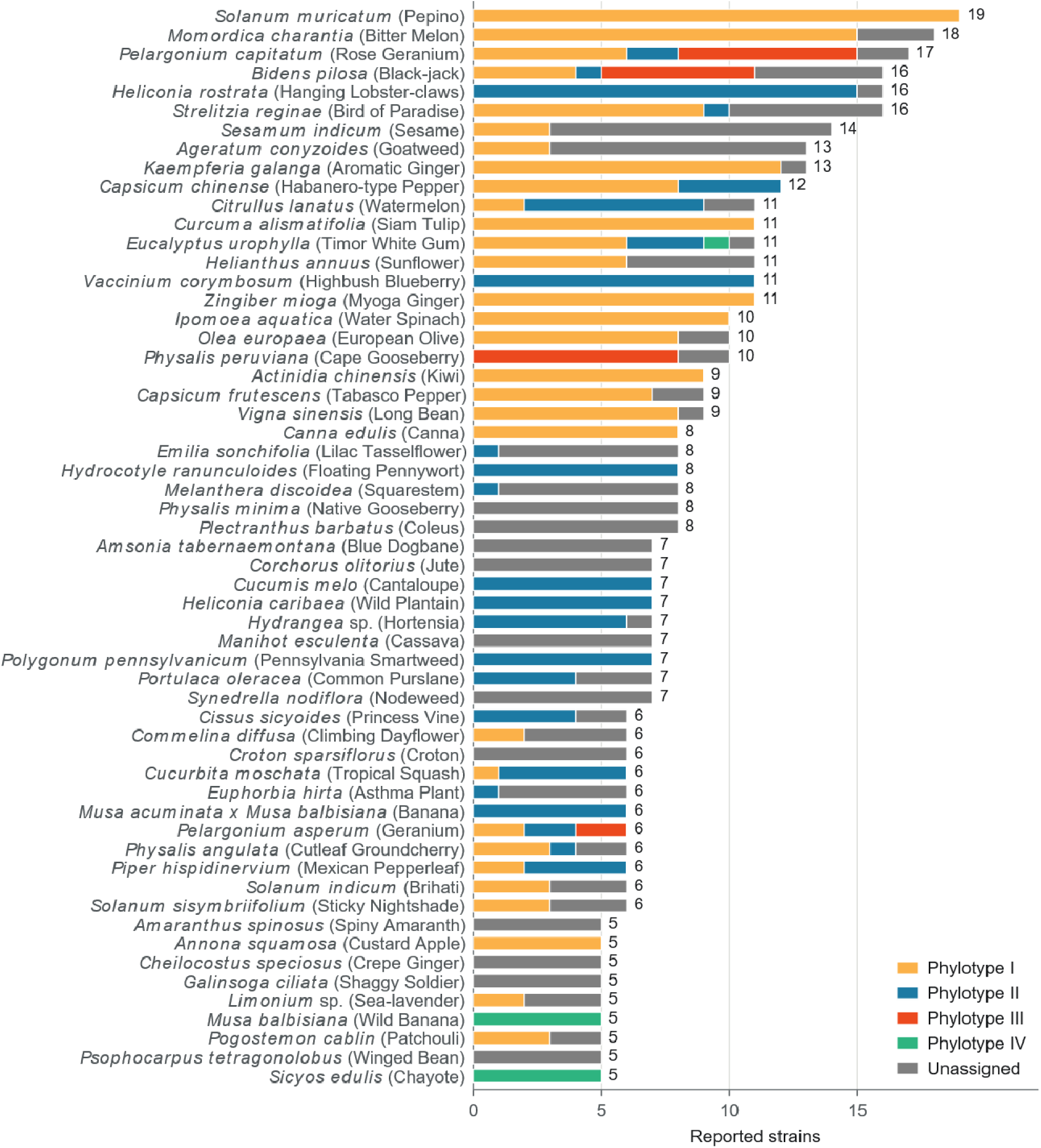
Phylotype composition of host species with 5 to 19 reported RSSC isolations. Bars give the number of reported strains of each phylotype, with the total printed at the right. This is a companion figure to Fig. 4.

### Biogeography

**Fig 6.**
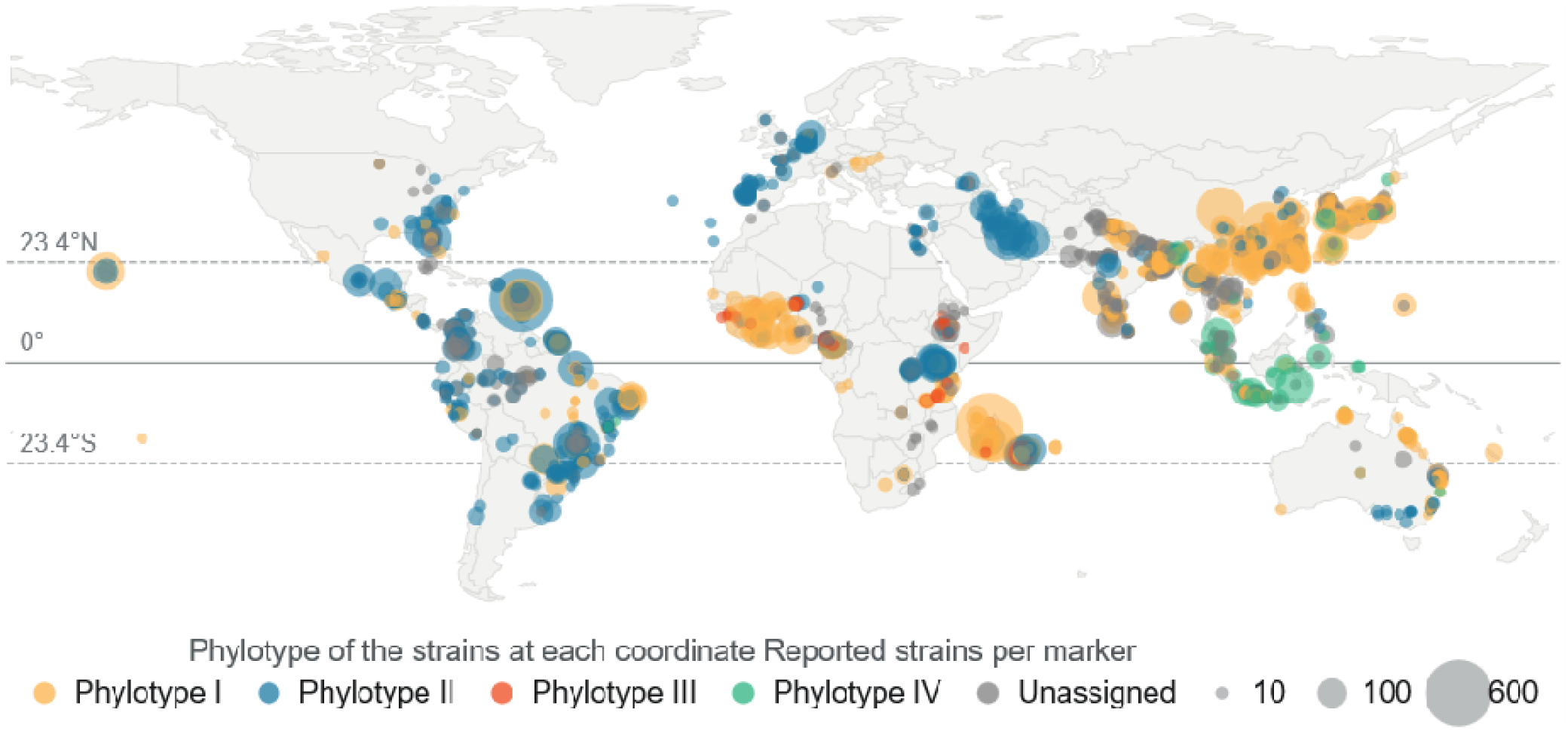
Phylotype distribution for RSSC strains reported at relatively specific locations. The figure plots 9,514 strains at 1,392 distinct coordinates as 1,732 circles. Each circle shows the number of strains isolated from specific coordinate locations, with area proportional to the number of reported strains. Circles were drawn in randomized order to reduce visual bias from overplotting. The additional strains recorded only to country level were not plotted here, but they are included in Fig 7.

**Fig 7.**
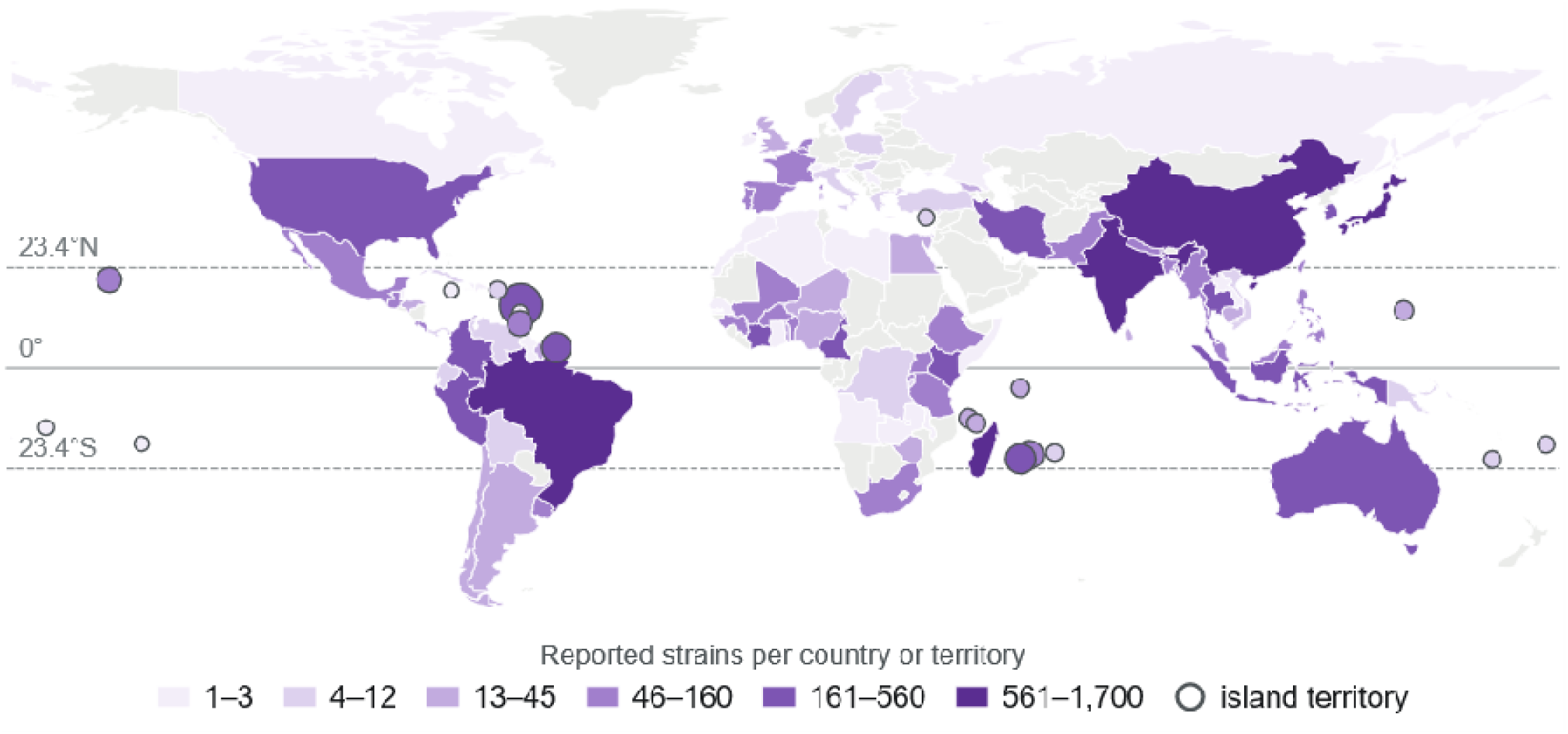
Reported distribution of the *Ralstonia solanacearum* species complex in the *Ralstonia* Diversity Dataset. Shading gives the number of reported strains per country or territory on a six-step logarithmic scale.

**Table 5:**
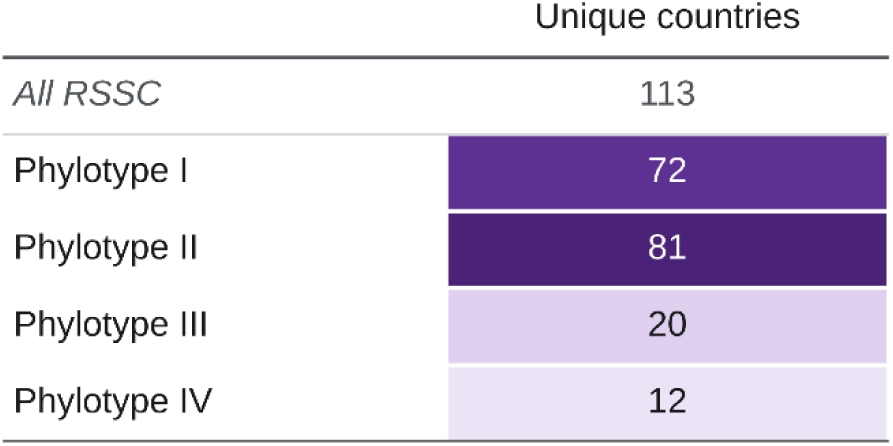
Relative distribution of the RSSC and each phylotype, based on the number of unique countries where the strain has been isolated.

### Biogeography and Host Range

**Fig 8.**
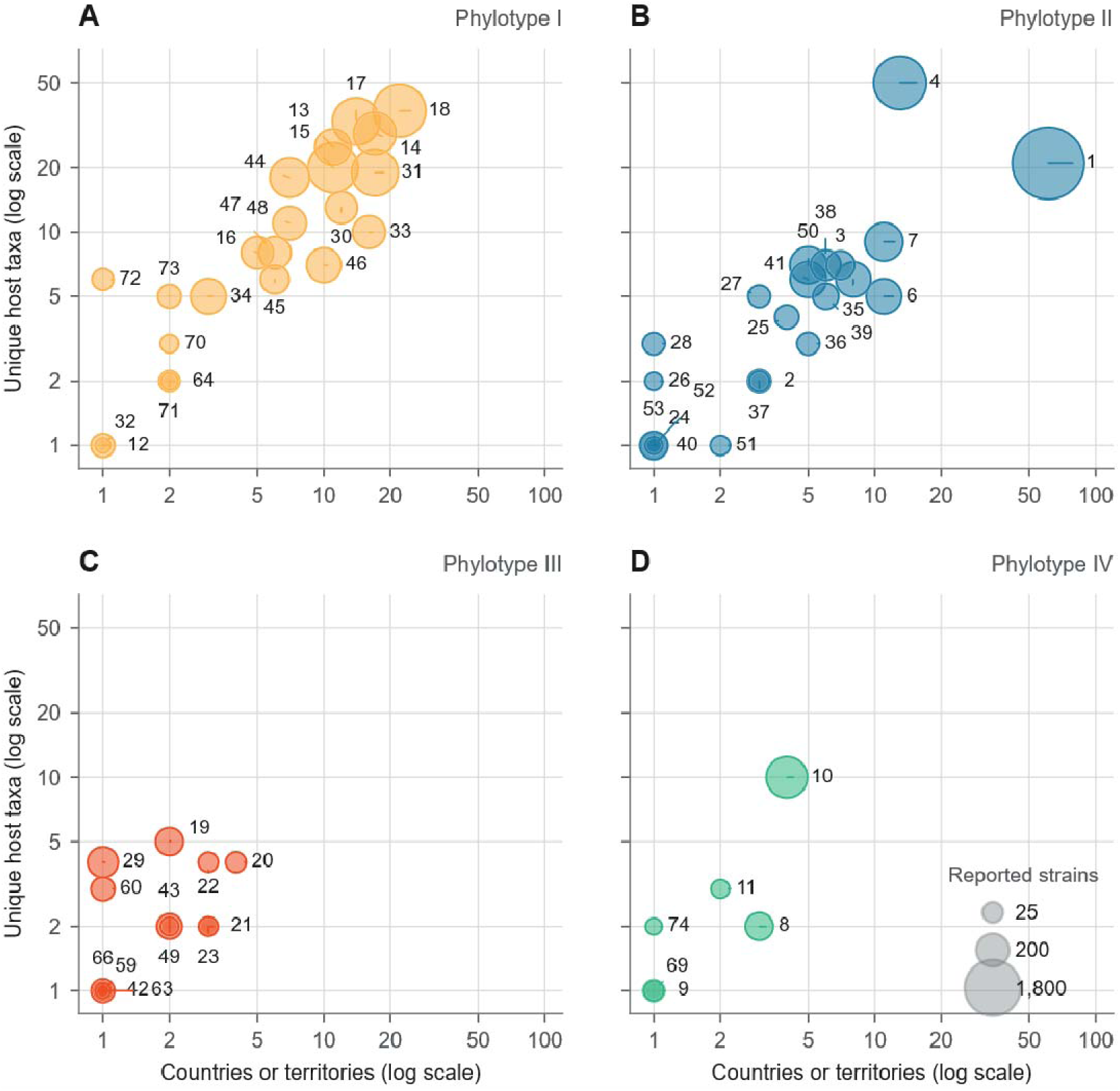
Geographic range against host range for the sequevars of each phylotype. (A) Phylotype I, (B) phylotype II, (C) phylotype III, (D) phylotype IV. Each circle is one sequevar, with area proportional to the number of reported strains on a scale common to all four panels; both axes are logarithmic and identical across panels. The 5 sequevars reported under more than one phylotype are omitted.

**Fig 9.**
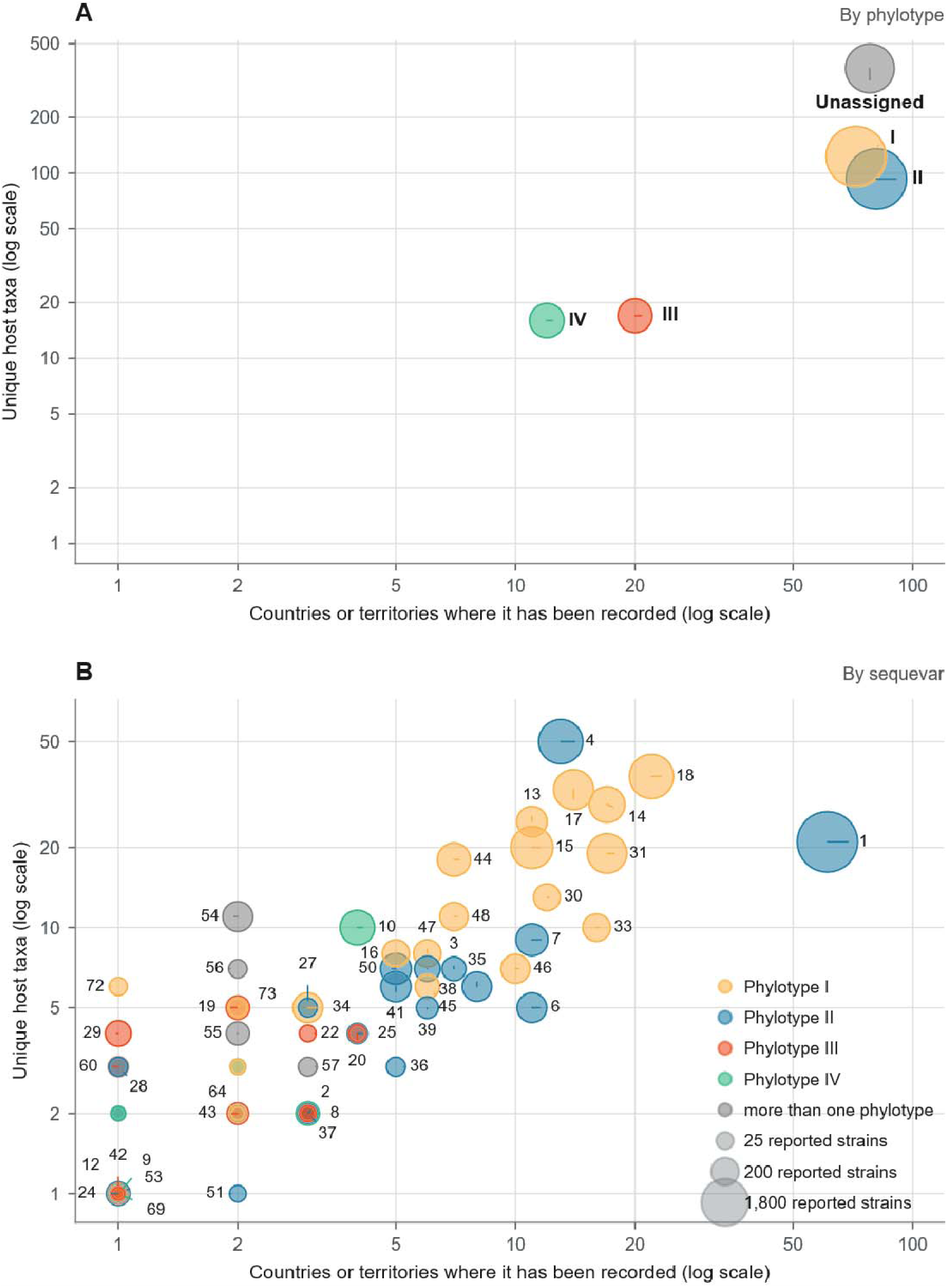
Geographic range against host range by phylotype. Circle area is proportional to the number of reported strains, and color gives the phylotype. Both axes are logarithmic. Circles are drawn at their exact positions, largest behind, so sequevars sharing a position appear as concentric rings; labels are offset and joined to their circle by a leader line of the same color, each placed nearer its own circle than any other. Sequevars with fewer than 6 reported strains are plotted without a label.

### RSSC Phylogenomics narrative on KBase

We created an open KBase narrative that has RSSC genomes labeled with a structured name that indicates their phylogenetic identity, strain name, host of isolation, subcontinent of isolation, and an NCBI accession number. The narrative is located at a new location because we decided to change the syntax of the genome names and because the prior narrative was slow to load and run. The prior narrative is listed in the previous version of this bioRxiv pre-print.

- Narrative with 825 public genomes: https://narrative.kbase.us/narrative/189849

Anyone can create a copy of these narratives, upload their own assembly to KBase, annotate the genes, and re-build the 49-gene phylogenetic tree to identify the position of new genomes. We will continue to update the narrative as we add genomes, so our colleagues should create a copy of the latest version of the narrative to assess new genomes.

**Fig 10.**
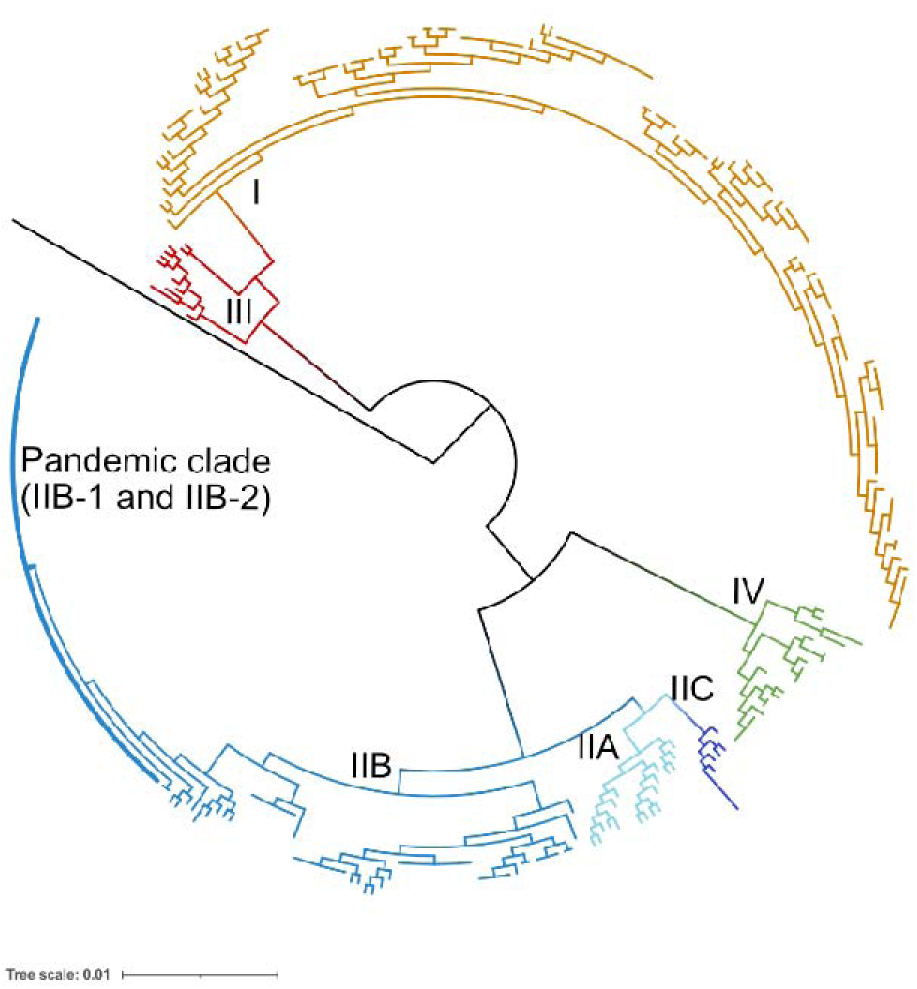
A phylogenetic tree of RSSC strains. The outgroup is an *R. mannitolilytica* genome. To create the stylized tree, the Newick format tree was downloaded from the output of the KBase SpeciesTree app, stylized in iTOL [357], and annotated in Affinity Designer. A high-resolution version (PDF, PNG, or Newick format) can be downloaded from the KBase narrative. The PDF has each genome labeled in searchable text.

### Assessing quality of genomes for downstream analyses

Genome assemblies vary in quality and can suffer from many flaws. We did not import genomes that NCBI flagged as “many frameshifted proteins” (often due to being a Nanopore-only assembly) or “fragmented assembly” (4700+ contigs, which suggests that the user uploaded only the coding sequences instead of the actual assembly).

For the assemblies imported into the KBase narrative, we used CheckM to assess their quality. CheckM yields estimates of the genomes’ completeness (are all of the expected single-copy genes present in the genome assembly?) and contamination (are there multiple homologs of the expected single-copy genes, suggesting that an unrelated genome has contaminated the assembly?). For completeness, we set a cut-off of requiring genomes to have no more than 4 missing “single copy genes”. This generally corresponds to a 99% complete cut-off. For contamination, we set a cut-off of 1%.

CheckM ran into an error when processing eight genomes, so these could not be assessed. Based o cut-offs, we classified 39 genomes as low-quality (24 genomes for low completeness and 15 genomes for high contamination). All of the draft genomes from Ailloud et al. 2015 have low completeness, likely because they were assembled from very short Illumina reads (2□×□50-nt paired). These genomes are estimated at 96-98% complete, which means the assemblies could be missing upwards of 200 genes. Two additional genomes have very low completeness: phyl. IIA-38 genome P816 was 81% complete and phyl. I genome VT0801 was 91% complete. After filtering for quality, there are only four phylotype IIA high-quality genomes available, making this the least represented major branch of *Ralstonia* diversity.

Because many *Ralstonia* isolates have been shared amongst scientists, either directly or via deposition into strain collections, there are cases where multiple assemblies are available for the same strain. In cases where this is obvious (strains share the same name and overlapping metadata), we selected the best assembly based on CheckM contamination/completeness. When those values were identical, we selected the genome with the best N50 or fewest contigs.

## Conclusion

Bacterial wilt pathogens in the *Ralstonia* species complex are high-impact global pathogens. We created a strain database that we will regularly update to document the distribution and host range of *Ralstonia*.

## Supporting information

Table S1: Ralstonia Diversity Dataset v6

## Acknowledgements

We thank all our colleagues who have carried out diversity surveys for *Ralstonia* and published genome sequences to NCBI. We thank Aracely Hernandez Rivas, Bhavana Karthik, Jia Yu Liou, Hana Mohammed, Darrell Sparks, and Ruxi Zhang for their contributions to the database.

## References

1. Ingel B, Caldwell D, Duong F, Parkinson DY, McCulloh KA, Iyer-Pascuzzi AS, et al. Revisiting the source of wilt symptoms: X-ray microcomputed tomography provides direct evidence that *Ralstonia* biomass clogs xylem vessels. BioRxiv. 2021. p. 2021.03.19.436187. doi:10.1101/2021.03.19.436187

2. Remenant B, Coupat-Goutaland B, Guidot A, Cellier G, Wicker E, Allen C, et al. Genomes of three tomato pathogens within the *Ralstonia solanacearum* species complex reveal significant evolutionary divergence. BMC Genomics. 2010;11: 379. doi:10.1186/1471-2164-11-379

3. Safni I, Cleenwerck I, De Vos P, Fegan M, Sly L, Kappler U. Polyphasic taxonomic revision of the *Ralstonia solanacearum* species complex: proposal to emend the descriptions of *R. solanacearum* and *R. syzygii* and reclassify current *R. syzygii* strains. Int J Syst Evol Microbiol. 2014;2014;64: 3087–103. doi:10.

4. Prior P, Ailloud F, Dalsing BL, Remenant B, Sanchez B, Allen C. Genomic and proteomic evidence supporting the division of the plant pathogen *Ralstonia solanacearum* into three species. BMC Genomics. 2016;17: 90. doi:10.1186/s12864-016-2413-z

5. Sharma P, Johnson MA, Mazloom R, Allen C, Heath LS, Lowe-Power TM, et al. Meta-analysis of the *Ralstonia solanacearum* species complex (RSSC) based on comparative evolutionary genomics and reverse ecology. Microb Genom. 2022;8: 000791. Available: https://www.ncbi.nlm.nih.gov/pubmed/35297758

6. Fegan M, Prior P. How complex is the *Ralstonia solanacearum* species complex. Bacterial wilt disease and the Ralstonia solanacearum species complex. APS press St. Paul; 2005. pp. 449–461. Available: https://www.researchgate.net/profile/Philippe_Prior/publication/37628297_How_Complex_is_the_Ralstonia_Solanacearum_Species_Complex/links/541047830cf2f2b29a4079bb.pdf

7. Elmgreen VN, Ramirez B, Sharma R, Vinatzer BA, Lowe-Power TM, Pierce-Ward NT. An interactive dashboard for global reports of the Ralstonia solanacearum species complex. 2025. p. 001126.v2. doi:10.1099/acmi.0.001126.v2

8. Arkin AP, Cottingham RW, Henry CS, Harris NL, Stevens RL, Maslov S, et al. KBase: The United States Department of Energy Systems Biology Knowledgebase. Nat Biotechnol. 2018;36: 566–569. doi:10.1038/nbt.4163

9. Berg LA. Weed hosts of the SFR strain of *Pseudomonas solanacearum*, causal organism of bacterial wilt of bananas. Phytopathology. 1971. Available: https://agris.fao.org/agris-search/search.do?recordID=US201302239782

10. Pegg KG, Moffett ML. Host range of the ginger strain of *Pseudomonas solanacearum* in Queensland. Aust J Exp Agric. 1971;11: 696–698. doi:10.1071/ea9710696

11. Power. Moko, a new bacterial disease on banana and plantain in Surinam. Surinaamse landbouw. 1976. Available: https://agris.fao.org/agris-search/search.do?recordID=US201302522553

12. Graham J, Lloyd AB. *Solanum cinereum*, a wild host of *Pseudomonas solanacearum* biotype II. Journal of the Australian Institute of Agricultural Science. 1978;44: 124–126. Available: https://www.cabdirect.org/cabdirect/abstract/19791354146

13. Kishun, Sohi, Rao. Two new collateral hosts for *Pseudomonas solanacearum*. Curr Sci. 1980. Available: https://www.cabdirect.org/cabdirect/abstract/19801370089

14. Velupillai, Stall. Variation among strains of *Pseudomonas solanacearum* from Florida. Proceedings of the Florida State. 1984. Available: https://journals.flvc.org/fshs/article/download/95368/91383

15. Hayward, AC. Bacterial wilt caused by *Pseudomonas solanacearum* in Asia and Australia: an overview. In: Persley GJ, editor. Bacterial Wilt Disease in Asia and the South Pacific. AICAR Conference Proceedings; 1986. pp. 15–24.

16. Iqbal, M, Kunmar, J. Bacterial wilt in Fiji. In: Persley GJ, editor. Bacterial Wilt Disease in Asia and the South Pacific. AICAR Conference Proceedings; 1986. pp. 15–24.

17. Sinha, SK. Bacterial wilt in India. In: Persley GJ, editor. Bacterial Wilt Disease in Asia and the South Pacific. AICAR Conference Proceedings; 1986. pp. 28–29.

18. Machmud, M. Bacterial wilt in Indonesia. In: Persley GJ, editor. Bacterial Wilt Disease in Asia and the South Pacific. AICAR Conference Proceedings; 1986. pp. 28–30.

19. Tomlinson, DL, Gunther, MT. Bacterial wilt in Papua New Guinea. In: Persley GJ, editor. Bacterial Wilt Disease in Asia and the South Pacific. AICAR Conference Proceedings; 1986. pp. 35–39.

20. He LY. Bacterial wilt in the People’s Republic of China. In: Persley GJ, editor. Bacterial Wilt Disease in Asia and the South Pacific. AICAR Conference Proceedings; 1986. pp. 35–39.

21. Valdez RB. Bacterial wilt in the Philippines. In: Persley GJ, editor. Bacterial Wilt Disease in Asia and the South Pacific. AICAR Conference Proceedings; 1986. pp. 49–56.

22. Velupillai M. Bacterial wilt in Sri Lanka. In: Persley GJ, editor. Bacterial Wilt Disease in Asia and the South Pacific. AICAR Conference Proceedings; 1986. pp. 57–64.

23. Titatarn V. Bacterial wilt in Thailand. In: Persley GJ, editor. Bacterial Wilt Disease in Asia and the South Pacific. AICAR Conference Proceedings; 1986. pp. 65–67.

24. Tung PH. Bacterial wilt in Vietnam. In: Persley GJ, editor. Bacterial Wilt Disease in Asia and the South Pacific. AICAR Conference Proceedings; 1986. pp. 68–70.

25. Buddenhagen. Bacterial wilt revisited. Bacterial wilt disease in Asia and the. 1985. Available: https://ageconsearch.umn.edu/record/134643/files/PR013.pdf#page=124

26. Li X, Dorsch T, Del Dot T, Sly LI, Stackebrandt E, Hayward AC. Phylogeny of biovars of *Pseudomonas solanacearum* based on sequencing of 16S rRNA. In: Hartman GL, Hayward AC, editors. Bacterial Wilt (Proceedings from an International Conference held in Taiwan). AICAR Conference Proceedings; 1993. pp. 93–95.

27. Alvarez AM, Berestecky J, Stiles JI, Ferreira SA, Benedict AA. Serological and molecular approaches to identification of *Pseudomonas solanacearum* strains from *Heliconia*. In: Hartman GL, Hayward AC, editors. Bacterial Wilt (Proceedings from an International Conference held in Taiwan). AICAR Conference Proceedings; 1993. pp. 62–69.

28. Marin JE, El-Nashaar HM. Pathogenicity of the new phenotypes of *Pseudomonas solanacearum* from Peru. In: Hartman GL, Hayward AC, editors. Bacterial Wilt (Proceedings from an International Conference held in Taiwan). AICAR Conference Proceedings; 1993. pp. 70–77.

29. Adhikari TB, Manandhar JB, Hartman GL. Characterisation of *Pseudomonas solanacearum* and evaluation of tomatoes in Nepal. In: Hartman, G. L. and Hayward, A. C, editor. Bacterial Wilt (Proceedings from an International Conference held in Taiwan). ACIAR Proceedings Series; 1993. pp. 132–137.

30. Hong NX, Mehan VK. Research on bacterial wilt of groundnut in Vietnam. In: Hartman GL, Hayward AC, editors. Bacterial Wilt (Proceedings from an International Conference held in Taiwan). AICAR Conference Proceedings; 1993. pp. 219–220.

31. Hamidah S, Lum K. Bacterial wilt of groundnuts in Malaysia. In: Hartman GL, Hayward AC, editors. Bacterial Wilt (Proceedings from an International Conference held in Taiwan). AICAR Conference Proceedings; 1993. pp. 225–227.

32. Wall GC, Sanchez JL. A biocontrol agent for *Pseudomonas solanacearum*. In: Hartman GL, Hayward AC, editors. Bacterial Wilt (Proceedings from an International Conference held in Taiwan). AICAR Conference Proceedings; 1993. pp. 320–321.

33. Fucikovsky LZ, Santos MO. Advance of bacterial wilt in bananas in Mexico. In: Hartman GL, Hayward AC, editors. Bacterial Wilt (Proceedings from an International Conference held in Taiwan). AICAR Conference Proceedings; 1993. pp. 341–342.

34. Girard JC, Nicole JF, Chiron JJ, Gaubiac AM, Huvier O, Oudard B, et al. Bacterial wilt due to *Pseudomonas solanacearum* in Reunion: General situation and current research. In: Hartman GL, Hayward AC, editors. Bacterial Wilt (Proceedings from an International Conference held in Taiwan). AICAR Conference Proceedings; 1993. pp. 343–347.

35. Sood AK, Singh BM. Prevalence of bacterial wilt of solanaceous vegetables in the mid-hill subhumid zone of Himachal Pradesh, India. In: Hartman GL, Hayward AC, editors. Bacterial Wilt (Proceedings from an International Conference held in Taiwan). AICAR Conference Proceedings; 1993. pp. 358–361.

36. Feng J, Zhang M, Bai X, Han B, Liu T, Fan M, et al. One specific DNA piece in *Pseudomonas solanacearum* affecting *Arachis hypogaea*. In: Hartman GL, Hayward AC, editors. Bacterial Wilt (Proceedings from an International Conference held in Taiwan). ACIAR Proceedings Series; 1993. pp. 245–252.

37. Ramesh CR, Bandyopadhyay AK. Bacterial wilt of tomato in Andaman and Nicobar Islands. In: Hartman GL, Hayward AC, editors. Bacterial Wilt (Proceedings from an International Conference held in Taiwan). AICAR Conference Proceedings; 1993. pp. 355–357.

38. Abdullah H. Bacterial wilt in Malaysia: Hosts, disease incidence and geographical distribution. In: Hartman GL, Hayward AC, editors. Bacterial Wilt (Proceedings from an International Conference held in Taiwan). AICAR Conference Proceedings; 1993. pp. 334–337.

39. Smith JJ, Offord LC, Holderness M, Saddler GS. Genetic diversity of *Burkholderia solanacearum* (synonym *Pseudomonas solanacearum)* race 3 in Kenya. Appl Environ Microbiol. 1995;61: 4263–4268. doi:10.1128/aem.61.12.4263-4268.1995

40. Fegan M, Taghavi M, Sly LI, Hayward AC. Phylogeny, diversity and molecular diagnostics of *Ralstonia solanacearum.* Bacterial Wilt Disease, Molecular and Ecological Aspects. 1998. pp. 19–33.

41. Mao GZ, He LY. Relationship of wild type strain motility and interaction with host plants in *Ralstonia solanacearum*. Bacterial Wilt Disease, Molecular and Ecological Aspects. 1998. pp. 184–191.

42. Nicole JF, Cheron JJ, Girard JC, Luisetti J. A tentative explanation of the distribution, on Reunion Island, of bacterial wilt caused by either biovar 2 or biovar 3 of *Ralstonia solanacearum*. Bacterial Wilt Disease, Molecular and Ecological Aspects. 1998. pp. 343–350.

43. Raymundo AK, Aves-ILagon Y, Denny TP. Analysis of genetic variation of a population of banana infecting strains of *Ralstonia solanacearum*. Bacterial Wilt Disease, Molecular and Ecological Aspects. 1998. pp. 56–60.

44. Robertson AE. Factors affecting the population of *Ralstonia solanacearum* in a naturally infested field planted to tobacco. Bacterial Wilt Disease, Molecular and Ecological Aspects. 1998. pp. 369–378.

45. Silveira EB, Michereff SJ, Mariano RLR. Epidemiology of tomato bacterial wilt in Agreste region of Pernambuco State, Brazil,. Bacterial Wilt Disease, Molecular and Ecological Aspects. 1998. pp. 358–363.

46. Smith JJ, Kibata GN, Murimi ZK, Lum KY, Fernandez-Northcote E, Offord LC, et al. Biogeographic Studies on *Ralstonia solanacearum* Race I and 3 by Genomic Fingerprinting. Bacterial Wilt Disease, Molecular and Ecological Aspects. 1998. pp. 50–55.

47. Stefanova M. Current Situation of Bacterial Wilt *(Ralstonia solanacearum* Smith) in Cuba. Bacterial Wilt Disease, Molecular and Ecological Aspects. 1998. pp. 364–368.

48. Hayward AC, Elphinstone IG, Caffier D, Lanse D, Stefani E, French ER, et al. Round Table on Bacterial Wilt (Brown Rot) of Potato. Bacterial Wilt Disease, Molecular and Ecological Aspects. 1998. pp. 420–430.

49. Tusiime G, Adipala E, Opio F, Bhagsari AS. Weeds as latent hosts of *Ralstonia solanacearum* in highland Uganda: Implications to development of an integrated control package for bacterial wilt. Bacterial Wilt Disease. Berlin, Heidelberg: Springer Berlin Heidelberg; 1998. pp. 413–419. doi:10.1007/978-3-662-03592-4_63

50. Urquhart L, Mienie NJJ, Steyn PL. The effect of temperature, storage period and inoculum concentration on symptom development and survival of *Ralstonia solanacearum* in inoculated tubers. Bacterial Wilt Disease. Berlin, Heidelberg: Springer Berlin Heidelberg; 1998. pp. 351–357. doi:10.1007/978-3-662-03592-4_53

51. Van Der Wolf JM, Bonants PJM, Smith JJ, Hagenaar M, Nijhuis E, Van Beckhoven JRCM, et al. Genetic Diversity of *Ralstonia solanacearum* Race 3 in Western Europe determined by AFLP, RC-PFGE and Rep-PCR. Bacterial Wilt Disease. Berlin, Heidelberg: Springer Berlin Heidelberg; 1998. pp. 44–49. doi:10.1007/978-3-662-03592-4_6

52. Boudazin G, Claire Le Roux A, Josi K, Labarre P, Jouan B. Design of division specific primers of *Ralstonia solanacearum* and application to the identification of European isolates. Eur J Plant Pathol. 1999;105: 373–380. doi:10.1023/A:1008763111230

53. Lin, Hsu, Tzeng. Weed hosts of Ralstonia solanacearum in Taiwan. Plant Protection Bulletin (Taipei). 1999. Available: https://www.cabdirect.org/cabdirect/abstract/20002301682

54. Wenneker M, Verdel MSW, Groeneveld RMW, Kempenaar C, van Beuningen AR, Janse JD. *Ralstonia (Pseudomonas) solanacearum* Race 3 (Biovar 2) in surface water and natural weed hosts: first report on stinging nettle *(Urtica dioica)*. Eur J Plant Pathol. 1999;105: 307–315. doi:10.1023/A:1008795417575

55. Horita M, Tsuchiya K. Comparative analysis of Japanese and foreign strains of *Ralstonia solanacearum* based on 16S ribosomal RNA gene sequences. J Gen Plant Pathol. 2000;66: 132–137. doi:10.1007/pl00012934

56. Dhital SP, Thaveechai N, Shrestha SK. Characteristics of *Ralstonia solanacearum* strains of potato wilt disease from Nepal and Thailand. Nepal Agric Res J. 2011; 42–47. doi:10.3126/narj.v4i0.4868

57. Coelho Netto RA, Noda H, Boher B. *Melanthera discoidea:* Um novo hospedeiro de *Ralstonia solanacearum*. Fitopatol Bras. 2001;26: 781–781. doi:10.1590/S0100-41582001000400020

58. Lee Yung-An, Fan Shu-Chung, Chiu Ling-Ya, Hsia Kuo-Chiang. Isolation of an insertion sequence from *Ralstonia solanacearum* race 1 and its potential use for strain characterization and detection. Appl Environ Microbiol. 2001;67: 3943–3950. doi:10.1128/AEM.67.9.3943-3950.2001

59. Pradhanang PM, Elphinstone JG, Fox RTV. Identification of crop and weed hosts of *Ralstonia solanacearum* biovar 2 in the hills of Nepal. Plant Pathol. 2000;49: 403–413. doi:10.1046/j.1365-3059.2000.00480.x

60. Williamson L, Nakaho K, Hudelson B, Allen C. *Ralstonia solanacearum* Race 3, Biovar 2 strains isolated from geranium are pathogenic on potato. Plant Dis. 2002;86: 987–991. doi:10.1094/PDIS.2002.86.9.987

61. Salanoubat M, Genin S, Artiguenave F, Gouzy J, Mangenot S, Arlat M, et al. Genome sequence of the plant pathogen *Ralstonia solanacearum*. Nature. 2002;415: 497–502. doi:10.1038/415497a

62. Coelho Netto RA, Pereira BG, Noda H, Boher B. Murcha bacteriana no estado do Amazonas, Brasil. Fitopatol Bras. 2004;29: 17–23. doi:10.1590/S0100-41582004000100004

63. Dittapongpitch V, Surat S. Detection of *Ralstonia solanacearum* in soil and weeds from commercial tomato fields using immunocapture and the polymerase chain reaction. J Phytopathol. 2003;151: 239–246. doi:10.1046/j.1439-0434.2003.00714.x

64. Belalcázar, Rosales. El “Moko” del plátano y banano y el rol de las plantas hospederas en su epidemología. Manejo convencional y. 2003. Available: https://books.google.com/books?hl=en&lr=&id=Oez07rNnVloC&oi=fnd&pg=PA159&dq=moko+belalcazar&ots=fZjKlvKEJU&sig=e5kUhJCWLa55oRFFmDDIC04JL2M

65. Kumar A, Sarma YR. Characterization of *Ralstonia solanacearum* causing bacterial wilt of ginger in India. Indian Phytopathol. 2004;57: 12–17.

66. Kumar A, Sarma YR, Anandaraj M. Evaluation of genetic diversity of *Ralstonia solanacearum* causing bacterial wilt of ginger using REP-PCR and PCR-RFLP. Current Science. 2004;87: 1555–1561. Available: https://www.jstor.org/stable/24109034

67. Caruso P, Palomo JL, Bertolini E, Alvarez B, López MM, Biosca EG. Seasonal variation of *Ralstonia solanacearum* biovar 2 populations in a Spanish river: recovery of stressed cells at low temperatures. Appl Environ Microbiol. 2005;71: 140–148. doi:10.1128/AEM.71.1.140-148.2005

68. Horita M, Tsuchiya K, Ooshiro A. Characteristics of *Ralstonia solanacearum* Biovar N2 Strains in Asia. J Phytopathol (1986). 2005;153: 209–213. doi:10.1111/j.1439-0434.2005.00954.x

69. Li QQ, Feng JX, Tang JL, Lin W, Duan CJ, Ye YF, et al. *Siraita grosvenorii* (Luo Han Guo; Cucurbitaceae) is a new host of *Ralstonia solanacearum* in China. Plant Pathol. 2005;54: 811–811. doi:10.1111/j.1365-3059.2005.01227.x

70. Prior P, Fegan M. Recent development in the phylogeny and classification of *Ralstonia solanacearum*. In: Ji JM, editor. Proceedings of the 1st International Symposium on Tomato Diseases. International Society for Horticultural Science (ISHS); 2005. pp. 127–136. doi:10.17660/ActaHortic.2005.695.14

71. Castillo JA, Greenberg JT. Evolutionary dynamics of *Ralstonia solanacearum*. Appl Environ Microbiol. 2007;73: 1225–1238. doi:10.1128/AEM.01253-06

72. Fouché-Weich J, Poussier S, Trigalet-Demery D, Berger D, Coutinho T. Molecular identification of some African strains of *Ralstonia solanacearum* from eucalypt and potato. J Gen Plant Pathol. 2006;72: 369–373. doi:10.1007/s10327-006-0307-7

73. Alfenas AC, Mafia RG, Sartório RC, Binoti DHB, Silva RR, Lau D, et al. *Ralstonia solanacearum* em viveiros clonais de eucalipto no Brasil. Fitopatol Bras. 2006;31: 357–366. doi:10.1590/s0100-41582006000400005

74. Carmeille A, Prior P, Kodja H, Chiroleu F, Luisetti J, Besse P. Evaluation of resistance to race 3, biovar 2 of *Ralstonia solanacearum* in tomato germplasm. J Phytopathol. 2006;154: 398–402. doi:10.1111/j.1439-0434.2006.01112.x

75. Ji P, Allen C, Sanchez-Perez A, Yao J, Elphinstone JG, Jones JB, et al. New diversity of *Ralstonia solanacearum* strains associated with vegetable and ornamental crops in Florida. Plant Dis. 2007;91: 195–203. doi:10.1094/PDIS-91-2-0195

76. Lewis Ivey ML, Gardener BBM, Opina N, Miller SA. Diversity of *Ralstonia solanacearum* infecting eggplant in the Philippines. Phytopathology. 2007;97: 1467–1475. doi:10.1094/PHYTO-97-11-1467

77. Wicker E, Grassart L, Coranson-Beaudu R, Mian D, Guilbaud C, Fegan M, et al. *Ralstonia solanacearum* strains from Martinique (French West Indies) exhibiting a new pathogenic potential. Appl Environ Microbiol. 2007;73: 6790–6801. doi:10.1128/AEM.00841-07

78. Black R, Seal S, Abubakar Z, Nono-Womdim R, Swai I. Wilt pathogens of Solanaceae in Tanzania: *Clavibacter michiganensis* subsp. *michiganensis*, *Pseudomonas corrugata*, and *Ralstonia solanacearum*. Plant Dis. 2007;83: 1070. doi:10.1094/PDIS.1999.83.11.1070A

79. Horita M, Tsuchiya K. Genetic Diversity of Japanese Strains of *Ralstonia solanacearum*. Phytopathology. 2007;91: 399–407. doi:10.1094/PHYTO.2001.91.4.399

80. Jeong Y, Kim J, Kang Y, Lee S, Hwang I. Genetic diversity and distribution of Korean isolates of *Ralstonia solanacearum*. Plant Dis. 2007;91: 1277–1287. doi:10.1094/PDIS-91-10-1277

81. Lemessa F, Zeller W. Isolation and characterisation of *Ralstonia solanacearum* strains from Solanaceae crops in Ethiopia. J Basic Microbiol. 2007;47: 40–49. doi:10.1002/jobm.200610199

82. Seo S-T, Park J-H, Han K-S, Cheong S-R, Lee S-D. Genetic diversity of *Ralstonia solanacearum* strains isolated from pepper and tomato plants in Korea. Research in Plant Disease. 2007;13: 24–29. doi:10.5423/RPD.2007.13.1.024

83. Yu Q, Alvarez AM, Moore PH, Zee F, Kim MS, de Silva A, et al. Molecular diversity of *Ralstonia solanacearum* isolated from ginger in Hawaii. Phytopathology. 2007;93: 1124–1130. doi:10.1094/PHYTO.2003.93.9.1124

84. Hong JC, Momol MT, Jones JB, Ji P, Olson SM, Allen C, et al. Detection of *Ralstonia solanacearum* in irrigation ponds and aquatic weeds associated with the ponds in North Florida. Plant Dis. 2008;92: 1674–1682. doi:10.1094/PDIS-92-12-1674

85. Sanchez Perez A, Mejia L, Fegan M, Allen C. Diversity and distribution of *Ralstonia solanacearum* strains in Guatemala and rare occurrence of tomato fruit infection. Plant Pathol. 2008. Available: https://bsppjournals.onlinelibrary.wiley.com/doi/abs/10.1111/j.1365-3059.2007.01769.x

86. Coutinho TA, Roux J, Riedel K-H, Terblanche J, Wingfield MJ. First report of bacterial wilt caused by *Ralstonia solanacearum* on eucalypts in South Africa. For Pathol. 2008;30: 205–210. doi:10.1046/j.1439-0329.2000.00205.x

87. Kagona. The incidence of bacterial wilt (Ralstonia solanacearum) in informal potato planting materials used by farmers in dedza and ntcheu districts of Malawi. M.S., Norwegian University of Life Sciences. 2008. Available: http://www.umb.no/statisk/noragric/publications/master/2008_justin_dickson_zayamba_kagona.pdf

88. Khakvar R, Sijam K, Wong MY, Radu S, Jones J, Thong KL. Genomic diversity of *Ralstonia solanacearum* strains isolated from banana farms in west Malaysia. Plant Pathol J. 2008;7: 162–167. doi:10.3923/ppj.2008.162.167

89. Obregón Barrios M, Rodríguez Gaviria PA, Morales Osorio JG, Salazar Yepes M. HOSPEDANTES DE *Ralstonia solanacearum* EN PLANTACIONES DE BANANO Y PLÁTANO EN COLOMBIA. Rev Fac Nac Agron Medellin. 2008;61: 4518–4526. Available: http://www.scielo.org.co/scielo.php?script=sci_abstract&pid=S0304-28472008000200006&lng=en&nrm=iso&tlng=es

90. Ustun N, Ozakman M, Karahan A. Outbreak of *Ralstonia solanacearum* biovar 2 causing brown rot on potato in the Aegean Region of Turkey. Plant Dis. 2008;92: 973. doi:10.1094/PDIS-92-6-0973B

91. Norman DJ, Zapata M, Gabriel DW, Duan YP, Yuen JMF, Mangravita-Novo A, et al. Genetic diversity and host range variation of *Ralstonia solanacearum* strains entering North America. Phytopathology. 2009;99: 1070–1077. doi:10.1094/PHYTO-99-9-1070

92. Nouri S, Bahar M, Fegan M. Diversity of *Ralstonia solanacearum* causing potato bacterial wilt in Iran and the first record of phylotype II/biovar 2T strains outside South America. Plant Pathol. 2009;58: 243–249. doi:10.1111/j.1365-3059.2008.01944.x

93. Toukam GMS, Cellier G, Wicker E, Guilbaud C, Kahane R, Allen C, et al. Broad diversity of *Ralstonia solanacearum* strains in Cameroon. Plant Dis. 2009;93: 1123–1130. doi:10.1094/PDIS-93-11-1123

94. Wicker E, Grassart L, Coranson-Beaudu R, Mian D, Prior P. Epidemiological evidence for the emergence of a new pathogenic variant of *Ralstonia solanacearumin* Martinique (French West Indies). Plant Pathol. 2009;58: 853–861. doi:10.1111/j.1365-3059.2009.02098.x

95. Guidot A, Elbaz M, Carrère S, Siri MI, Pianzzola MJ, Prior P, et al. Specific genes from the potato brown rot strains of *Ralstonia solanacearum* and their potential use for strain detection. Phytopathology. 2009;99: 1105–1112. doi:10.1094/PHYTO-99-9-1105

96. Liu Y, Kanda A, Yano K, Kiba A, Hikichi Y, Aino M, et al. Molecular typing of Japanese strains of *Ralstonia solanacearum* in relation to the ability to induce a hypersensitive reaction in tobacco. J Gen Plant Pathol. 2009;75: 369–380. doi:10.1007/s10327-009-0188-7

97. Ramírez M, Moncada RN, Villegas-Escobar V, Jackson RW, Ramírez CA. Phylogenetic and pathogenic variability of strains of *Ralstonia solanacearum* causing moko disease in Colombia. Plant Pathol. 2019;69: 360–369. doi:10.1111/ppa.13121

98. Xu J, Pan ZC, Prior P, Xu JS, Zhang Z, Zhang H, et al. Genetic diversity of *Ralstonia solanacearum* strains from China. Eur J Plant Pathol. 2009;125: 641–653. doi:10.1007/s10658-009-9512-5

99. de Andrade FWR, da Rocha Amorim EP, Eloy AP, Rufino MJ. Ocorrência de Doenças em Bananeiras no Estado de Alagoas. Summa Phytopathologica. 2009;35: 305–309. doi:10.1590/S0100-54052009000400008

100. Cardozo C, Rodríguez P, Marín M. Molecular characterization of the *Ralstonia solanacearum* species complex in the banana growing region of Uraba. Agron Colombiana. 2009;27: 203–210. Available: http://www.scielo.org.co/scielo.php?pid=S0120-99652009000200008&script=sci_abstract&tlng=en

101. Cellier G, Prior P. Deciphering phenotypic diversity of *Ralstonia solanacearum* strains pathogenic to potato. Phytopathology. 2010;100: 1250–1261. doi:10.1094/PHYTO-02-10-0059

102. Horita M, Suga Y, Ooshiro A, Tsuchiya K. Analysis of genetic and biological characters of Japanese potato strains of *Ralstonia solanacearum*. J Gen Plant Pathol. 2010;76: 196–207. doi:10.1007/s10327-010-0229-2

103. Li J-G, Liu H-X, Cao J, Chen L-F, Gu C, Allen C, et al. PopW of *Ralstonia solanacearum*, a new two-domain harpin targeting the plant cell wall. Mol Plant Pathol. 2010;11: 371–381. doi:10.1111/j.1364-3703.2010.00610.x

104. Khoodoo MHR, Ganoo ES, Saumtally AS. Molecular characterization and epidemiology of *Ralstonia solanacearum* Race 3 biovar 2 Causing Brown Rot of Potato in Mauritius. J Phytopathol. 2010;158: 503–512. doi:10.1111/j.1439-0434.2009.01654.x

105. Singh D, Sinha S, Yadav DK, Sharma JP, Srivastava DK, Lal HC, et al. Characterization of biovar/races of *Ralstonia solanacearum*, the incitant of bacterial wilt in solanaceous crops. Indian Phytopath. 2010;63: 261–265.

106. Stevens P, van Elsas JD. Genetic and phenotypic diversity of *Ralstonia solanacearum* biovar 2 strains obtained from Dutch waterways. Antonie Van Leeuwenhoek. 2010;97: 171–188. doi:10.1007/s10482-009-9400-1

107. Chandrashekara KN, Prasannakumar MK. New host plants for *Ralstonia solanacearum* from India. Plant Pathol. 2010;59: 1164–1164. doi:10.1111/j.1365-3059.2010.02358.x

108. Thera AT, Jacobsen BJ, Neher OT. Bacterial Wilt of Solanaceae Caused by *Ralstonia solanacearum* Race 1 Biovar 3 in Mali. Plant Dis. 2010;94: 372. doi:10.1094/PDIS-94-3-0372B

109. Kubota R, Schell MA, Peckham GD, Rue J, Alvarez AM, Allen C, et al. In silico genomic subtraction guides development of highly accurate, DNA-based diagnostics for *Ralstonia solanacearum* race 3 biovar 2 and blood disease bacterium. J Gen Plant Pathol. 2011;77: 182–193. doi:10.1007/s10327-011-0305-2

110. Lebeau A, Daunay M-C, Frary A, Palloix A, Wang J-F, Dintinger J, et al. Bacterial wilt resistance in tomato, pepper, and eggplant: genetic resources respond to diverse strains in the *Ralstonia solanacearum* species complex. Phytopathology. 2011;101: 154–165. doi:10.1094/PHYTO-02-10-0048

111. Pinheiro CR, Amorim JAE, Diniz LEC, Silva AMF da, Talamini V, Souza Júnior MT. Diversidade genética de isolados de *Ralstonia solanacearum* e caracterização molecular quanto a filotipos e sequevares. Pesqui Agropecu Bras. 2011;46: 593–602. doi:10.1590/S0100-204X2011000600004

112. Remenant B, de Cambiaire J-C, Cellier G, Jacobs JM, Mangenot S, Barbe V, et al. *Ralstonia syzygii*, the Blood Disease Bacterium and some Asian *R. solanacearum* strains form a single genomic species despite divergent lifestyles. PLoS One. 2011;6: e24356. doi:10.1371/journal.pone.0024356

113. Siri MI, Sanabria A, Pianzzola MJ. Genetic diversity and aggressiveness of *Ralstonia solanacearum* strains causing bacterial wilt of potato in Uruguay. Plant Dis. 2011;95: 1292–1301. doi:10.1094/PDIS-09-10-0626

114. Xue Q-Y, Yin Y-N, Yang W, Heuer H, Prior P, Guo J-H, et al. Genetic diversity of *Ralstonia solanacearum* strains from China assessed by PCR-based fingerprints to unravel host plant- and site-dependent distribution patterns. FEMS Microbiol Ecol. 2011;75: 507–519. doi:10.1111/j.1574-6941.2010.01026.x

115. Li Z, Wu S, Bai X, Liu Y, Lu J, Liu Y, et al. Genome sequence of the tobacco bacterial wilt pathogen *Ralstonia solanacearum*. J Bacteriol. 2011;193: 6088–6089. doi:10.1128/JB.06009-11

116. Chaudhry, Rashid. Isolation and characterization of *Ralstonia solanacearum* from infected tomato plants of Soan Skesar valley of Punjab. Pak J Bot. 2011. Available: https://www.academia.edu/download/61150112/55_Potato20191106-37160-1z0yurr.pdf

117. Mondal, Bhattacharya, Khatua. Crop and weed host of *Ralstonia solanacearum* in West Bengal. Journal of Crop and Weed. 2011. Available: https://www.cabi.org/ISC/FullTextPDF/2014/20143063677.pdf

118. Izadiyan M, Taghavi S. Diversity of Iranian isolates of *Ralstonia solanacearum*. Phytopathologia Mediterranea. 2011;50: 236–244. doi:10.2307/26458697

119. Adebayo OS, Nwanguma EI. Incidence of *Phytophthora* blight and bacterial wilt in dry season pepper production in Katsina, Nigeria. Acta Hortic. 2011;917. Available: https://www.actahort.org/books/917/917_22.htm

120. Bekele B, Abate E, Asefa A, Dickinson M. Incidence of potato viruses and bacterial wilt disease in the west Amhara sub-region of Ethiopia. Journal of Plant Pathology. 2011;93: 149–157. doi:10.4454/JPP.V93I1.285

121. Chandrashekara K, Prasannakumar M, Deepa M, Vani A. Coleus, a new host for *Ralstonia solanacearum* race 1 biovar 3 in India. Journal of Plant Pathology. 2011;93: 233–235. doi:10.4454/JPP.V93I1.298

122. Liu Q, Li Y, Chen J. First Report of Bacterial Wilt Caused by *Ralstonia solanacearum* on *Mesona chinensis* in China. Plant Dis. 2011;95: 222. doi:10.1094/PDIS-08-10-0603

123. Ruhl G, Twieg E, DeVries R, Levy L, Byrne J, Mollov D, et al. First report of bacterial wilt in *Mandevilla* (= *Dipladenia*) *splendens* “Red Riding Hood” in the United States caused by *Ralstonia solanacearum* biovar 3. Plant Dis. 2011;95: 614. doi:10.1094/PDIS-11-10-0858

124. Chen Y-J, Lin Y-S, Chung W-H. Bacterial wilt of sweet potato caused by *Ralstonia solanacearum* in Taiwan. J Gen Plant Pathol. 2012;78: 80–84. doi:10.1007/s10327-011-0353-7

125. Cellier G, Remenant B, Chiroleu F, Lefeuvre P, Prior P. Phylogeny and population structure of brown rot- and Moko disease-causing strains of *Ralstonia solanacearum* phylotype II. Appl Environ Microbiol. 2012;78: 2367–2375. doi:10.1128/AEM.06123-11

126. N’guessan CA, Abo K, Fondio L, Chiroleu F, Lebeau A, Poussier S, et al. So near and yet so far: the specific case of *Ralstonia solanacearum* populations from Côte d’Ivoire in Africa. Phytopathology. 2012;102: 733–740. doi:10.1094/PHYTO-11-11-0300

127. Hong JC, Norman DJ, Reed DL, Momol MT, Jones JB. Diversity among *Ralstonia solanacearum* strains isolated from the Southeastern United States. Phytopathology. 2012;102: 924–936. doi:10.1094/phyto-12-11-0342

128. Wairuri CK, van der Waals JE, van Schalkwyk A, Theron J. *Ralstonia solanacearum* needs Flp pili for virulence on potato. Mol Plant Microbe Interact. 2012;25: 546–556. doi:10.1094/MPMI-06-11-0166

129. Mondal B, Mahapatra S, Khatua DC. Record of some new diseases of horticultural plants of West Bengal. Journal of Interacademicia. 2012;16: 34–43. Available: https://www.researchgate.net/profile/Sunita-Mahapatra/publication/307907016_Records_of_some_new_diseases_of_horticultural_plants_of_West_Bengal/links/5dd52315299bf11ec8630ad2/Records-of-some-new-diseases-of-horticultural-plants-of-West-Bengal.pdf

130. Begum N, Haque MI, Mukhtar T, Naqvi SM, Wan JF. Status of bacterial wilt caused by *Ralstonia solanacearum* in Pakistan. Pakistan journal of phytopathology. 2012;24: 11–20. Available:

131. Chandrashekara K, Prasannakumar M, Deepa M, Vani A, Khan A. Prevalence of Races and biotypes of *Ralstonia solanacearum* in India. J Plant Prot Res. 2012;52: 53–58. doi:10.2478/v10045-012-0009-4

132. Mepharishvili G, Sikharulidze Z, Thwaites R, Tsetskhladze T, Dumbadze R, Gabaidze M, et al. First confirmed report of bacterial wilt of tomato in Georgia caused by *Ralstonia solanacearum*. New Dis Rep. 2012;25: 16–16. doi:10.5197/j.2044-0588.2012.025.016

133. Huang Q, Yan X, Wang J-F. Improved biovar test for *Ralstonia solanacearum*. J Microbiol Methods. 2012;88: 271–274. doi:10.1016/j.mimet.2011.12.007

134. Cruz L, Eloy M, Quirino F, Silva HO da, Tenreiro R. Molecular epidemiology of *Ralstonia solanacearum* strains from plants and environmental sources in Portugal. Eur J Plant Pathol. 2012;133: 687–706. doi:10.1007/s10658-012-9947-y

135. Ramsubhag A, Lawrence D, Cassie D, Fraser R, Umaharan P, Prior P, et al. Wide genetic diversity of *Ralstonia solanacearum* strains affecting tomato in Trinidad, West Indies. Plant Pathol. 2012;61: 844–857. doi:10.1111/j.1365-3059.2011.02572.x

136. Rodrigues LMR, Destéfano SAL, da Silva MJ, Costa GGL, Maringoni AC. Characterization of Ralstonia solanacearum strains from Brazil using molecular methods and pathogenicity tests. J Plant Pathol. 2012;94: 505–516. Available: http://www.jstor.org/stable/45156277

137. Prieto Romo J, Morales Osorio JG, Salazar Yepes M. Identification of new hosts for *Ralstonia solanacearum* (Smith) race 2 from Colombia. Rev Prot Veg. 2012;27: 151–161. Available: http://scielo.sld.cu/scielo.php?script=sci_abstract&pid=S1010-27522012000300003&lng=en&nrm=iso&tlng=en

138. Sumangala K, Lingaraju S, Hegde Y, Byadagi AS. Genetic diversity of *Ralstonia solanacaerum* from major tomato growing areas of Karnataka. International Journal of Plant Protection. 2012;5: 324–328. Available: https://www.semanticscholar.org/paper/Genetic-diversity-of-Ralstonia-solanacaerum-from-of-Sumangala-Lingaraju/7f18b566f4bf578f4bab86667a94bd99f2487da9

139. Santana BG, Lopes CA, Alvarez E, Barreto CC, Allen C, Quirino BF. Diversity of Brazilian biovar 2 strains of *Ralstonia solanacearum*. J Gen Plant Pathol. 2012;78: 190–200. doi:10.1007/s10327-012-0369-7

140. Cao Y, Tian B, Liu Y, Cai L, Wang H, Lu N, et al. Genome sequencing of *Ralstonia solanacearum* FQY_4, isolated from a bacterial wilt nursery used for breeding crop resistance. Genome Announc. 2013;1. doi:10.1128/genomeA.00125-13

141. Garcia AL, Lima WG, Souza EB, Michereff SJ, Mariano RLR. Characterization of *Ralstonia solanacearum* causing bacterial wilt in bell pepper in the state of Pernambuco, Brazil. J Plant Pathol. 2013;95: 237–245. Available: http://www.jstor.org/stable/23721514

142. Izadiyan M, Taghavi SM. Host range variation and genetic diversity of Iranian isolates of *Ralstonia solanacearum* from potato and tomato with RAPD and (GTG)5-PCR. J Plant Pathol. 2013;95: 87–97. Available: http://www.jstor.org/stable/23721740

143. N’guessan CA, Brisse S, Le Roux-Nio A-C, Poussier S, Koné D, Wicker E. Development of variable number of tandem repeats typing schemes for *Ralstonia solanacearum*, the agent of bacterial wilt, banana Moko disease and potato brown rot. J Microbiol Methods. 2013;92: 366–374. doi:10.1016/j.mimet.2013.01.012

144. Pan ZC, Xu J, Prior P, Xu JS, Zhang H, Chen KY, et al. Development of a specific molecular tool for the detection of epidemiologically active mulberry causing-disease strains of *Ralstonia solanacearum* phylotype I (historically race 5-biovar 5) in China. Eur J Plant Pathol. 2013;137: 377–391. Available: https://idp.springer.com/authorize/casa?redirect_uri=https://link.springer.com/article/10.1007/s10658-013-0249-9&casa_token=SRqxC6fKvn0AAAAA:W2Y0yW7TOoI6xE3C-2W0ldQZ3IYr-2ZbkPM7hFrWg-k9GQz8lJD5FvXlknp-K7v9DIrkhAMv0l-_XRzu

145. Parkinson N, Bryant R, Bew J, Conyers C, Stones R, Alcock M, et al. Application of variable-number tandem-repeat typing to discriminate *Ralstonia solanacearum* strains associated with English watercourses and disease outbreaks. Appl Environ Microbiol. 2013;79: 6016–6022. doi:10.1128/AEM.01219-13

146. Waki T, Horita M, Kurose D, Mulya K, Tsuchiya K. Genetic Diversity of Zingiberaceae Plant Isolates of *Ralstonia solanacearum* in the Asia-Pacific Region. Japan Agricultural Research Quarterly: JARQ. 2013;47: 283–294. doi:10.6090/jarq.47.283

147. Ahmed N, Islam M, Hossain M, Meah M, Hossain M. Determination of races and biovars of *Ralstonia solanacearum* causing bacterial wilt disease of potato. J Agric Sci. 2013;5: 86–93. doi:10.5539/jas.v5n6p86

148. Kumar R, Barman A, Jha G, Ray SK. Identification and establishment of genomic identity of *Ralstonia solanacearum* isolated from a wilted chilli plant at Tezpur, North East India. Curr Sci. 2013;105: 1571–1578. Available: http://www.jstor.org/stable/24098856

149. Pradhanang PM, Momol MT. Survival of *Ralstonia solanacearum* in soil under irrigated rice culture and aquatic weeds. J Phytopathol. 2013;149: 707–711. doi:10.1046/j.1439-0434.2001.00700.x

150. Milijašević-Marčić S, Todorovic B, Potočnik I, Rekanović E, Stepanovic M, Mitrović J, et al. *Ralstonia solanacearum* - a New Threat to Potato Production in Serbia. Pesticidi I Fitomedicina. 2013;28: 229–237. doi:10.2298/PIF1304229M

151. Romero GC, de Jensen CE, Palmateer AJ. First report of tomato wilt caused by *Ralstonia solanacearum* Biovar 1 in Puerto Rico. Plant Health Prog. 2013;14: 53. doi:10.1094/PHP-2013-0418-01-BR

152. Shan W, Yang X, Ma W, Yang Y, Guo X, Guo J, et al. Draft genome sequence of *Ralstonia solanacearum* race 4 biovar 4 strain SD54. Genome Announc. 2013;1. doi:10.1128/genomeA.00890-13

153. She X-M, He Z-F, Luo F-F, Li H-P. First Report of Bacterial Wilt Caused by *Ralstonia solanacearum* on *Ageratum conyzoides* in China. Plant Dis. 2013;97: 418. doi:10.1094/PDIS-08-12-0780-PDN

154. Avinash P, Umesha S. Identification and genetic diversity of bacterial wilt pathogen in brinjal. Arch Phytopathol Pflanzenschutz. 2014;47: 398–406. doi:10.1080/03235408.2013.811017

155. Wu Y-F, Cheng A-S, Lin C-H, Chen C-Y. First Report of Bacterial Wilt Caused by *Ralstonia solanacearum* on Roselle in Taiwan. Plant Dis. 2013;97: 1375. doi:10.1094/PDIS-02-13-0186-PDN

156. Albuquerque GMR, Santos LA, Felix KCS, Rollemberg CL, Silva AMF, Souza EB, et al. Moko disease-causing strains of *Ralstonia solanacearum* from Brazil extend known diversity in paraphyletic Phylotype II. Phytopathology. 2014;104: 1175–1182. doi:10.1094/PHYTO-12-13-0334-R

157. Lin C-H, Tsai K-C, Prior P, Wang J-F. Phylogenetic relationships and population structure of *Ralstonia solanacearum* isolated from diverse origins in Taiwan. Plant Pathol. 2014;63: 1395–1403. doi:10.1111/ppa.12209

158. Fonseca NR, Guimarães LMS, Hermenegildo PS, Teixeira RU, Lopes CA, Alfenas AC. Molecular characterization of *Ralstonia solanacearum* infecting *Eucalyptus* spp. in Brazil. For Pathol. 2014;44: 107–116. doi:10.1111/efp.12073

159. Horita M, Tsuchiya K, Suga Y, Yano K, Waki T, Kurose D, et al. Current classification of *Ralstonia solanacearum* and genetic diversity of the strains in Japan. J Gen Plant Pathol. 2014;80: 455–465. doi:10.1007/s10327-014-0537-z

160. Kumar A, Prameela TP, Panja B. Genetic characterization of an Indian isolate of *Ralstonia solanacearum* race 3/ biovar 2/ phylotype IIB from potato. Indian Phytopath. 2014;67: 346–352.

161. Ramesh R, Achari GA, Gaitonde S. Genetic diversity of *Ralstonia solanacearum* infecting solanaceous vegetables from India reveals the existence of unknown or newer sequevars of Phylotype I strains. Eur J Plant Pathol. 2014;140: 543–562. doi:10.1007/s10658-014-0487-5

162. Sagar V, Jeevalatha A, Mian S, Chakrabarti SK, Gurjar MS, Arora RK, et al. Potato bacterial wilt in India caused by strains of phylotype I, II and IV of *Ralstonia solanacearum*. Eur J Plant Pathol. 2014;138: 51–65. doi:10.1007/s10658-013-0299-z

163. Sujeeun L. Rôle des effecteurs de type III dans l’interaction *Ralstonia solanacearum/* aubergine: approches de génétique évolutive et fonctionnelle. Université de la Réunion. 2014. Available: https://agritrop.cirad.fr/573610/

164. Waki T, Horita M. Grouping of *Ralstonia solanacearum* strains in Tochigi Prefecture based on pathogenicity to *Solanum* plants and discriminating pathogenicity groups by PCR. Jpn J Phytopathol. 2014;80: 229–234. doi:10.3186/jjphytopath.80.229

165. Zulperi D, Sijam K, Ahmad ZAM, Awang Y, Mohd Hata E. Phylotype classification of *Ralstonia solanacearum* biovar 1 strains isolated from banana *(Musa* spp.) in Malaysia. Archives of Phytopathology and Plant Protection. 2014;47: 2352–2364. doi:10.1080/03235408.2013.876748

166. Deberdt P, Guyot J, Coranson-Beaudu R, Launay J, Noreskal M, Rivière P, et al. Diversity of *Ralstonia solanacearum* in French Guiana expands knowledge of the “Emerging Ecotype.” Phytopathology. 2014;104: 586–596. doi:10.1094/PHYTO-09-13-0264-R

167. Li X, Nie J, Hammill DL, Smith D, Xu H, De Boer SH. A comprehensive comparison of assays for detection and identification of *Ralstonia solanacearum* race 3 biovar 2. J Appl Microbiol. 2014;117: 1132–1143. doi:10.1111/jam.12585

168. Santiago TR, Grabowski C, Mizubuti ESG. First report of bacterial wilt caused by *Ralstonia solanacearum* on *Eucalyptus* sp. in Paraguay. New Dis Rep. 2014;29: 2–2. doi:10.5197/j.2044-0588.2014.029.002

169. Shrestha G, Prajapati S, Mahato BN. Plant diseases and their management practices in commercial organic and conventional vegetable farms in Kathmandu valley. Nepalese Journal of Agricultural Sciences. 2014;12: 129–141. Available: https://d1wqtxts1xzle7.cloudfront.net/83861157/Plant_diseases_and_their_management_prac20220411-5949-zyq7pt.pdf

170. Subedi N, Gilbertson RL, Osei MK, Cornelius E, Miller SA. First report of bacterial wilt caused by *Ralstonia solanacearum* in Ghana, west Africa. Plant Dis. 2014;98: 840. doi:10.1094/PDIS-09-13-0963-PDN

171. Tebaldi ND, Leite LN, Marque JM de, Furlanetto MCA, Mota LCBM. Occurrence of *Ralstonia solanacearum* on olive tree in Brazil. Summa Phytopathol. 2014;40: 185–185. doi:10.1590/0100-5405/1983

172. Ajitomi A, Inoue Y, Horita M, Nakaho K. Bacterial wilt of three *Curcuma* species, *C. longa* (turmeric), *C. aromatica* (wild turmeric) and *C. zedoaria* (zedoary) caused by *Ralstonia solanacearum* in Japan. J Gen Plant Pathol. 2015;81: 315–319. doi:10.1007/s10327-015-0596-9

173. Albuquerque P, Marcal ARS, Caridade C, Costa R, Mendes MV, Tavares F. A quantitative hybridization approach using 17 DNA markers for identification and clustering analysis of *Ralstonia solanacearum*. Plant Pathol. 2015;64: 1270–1283. doi:10.1111/ppa.12386

174. Ayin CM, Schlub RL, Yasuhara-Bell J, Alvarez AM. Identification and characterization of bacteria associated with decline of ironwood *(Casuarina equisetifolia)* in Guam. Australas Plant Pathol. 2015;44: 225–234. doi:10.1007/s13313-014-0341-4

175. Cellier G, Moreau A, Chabirand A, Hostachy B, Ailloud F, Prior P. A duplex PCR assay for the detection of *Ralstonia solanacearum* phylotype II strains in *Musa* spp. PLoS One. 2015;10: e0122182. doi:10.1371/journal.pone.0122182

176. Clarke CR, Studholme DJ, Hayes B, Runde B, Weisberg A, Cai R, et al. Genome-enabled phylogeographic investigation of the quarantine pathogen *Ralstonia solanacearum* Race 3 Biovar 2 and screening for sources of resistance against its core effectors. Phytopathology. 2015;105: 597–607. doi:10.1094/PHYTO-12-14-0373-R

177. Naranjo Feliciano E, Yglesia Lozano A, García A, Martínez Zubiaur Y. Diversidad genética de aislados de *Ralstonia solanacearum* de Cuba, mediante amplificación de las regiones repetitivas del genoma (REP-PCR). Rev Protección Veg. 2015;30: 52–59. Available: http://scielo.sld.cu/scielo.php?pid=S1010-27522015000100009&script=sci_arttext&tlng=pt

178. Gurjar MS, Sagar V, Bag TK, Singh BP, Sharma S, Jeevalatha A, et al. Genetic diversity of *Ralstonia solanacearum* strains causing bacterial wilt of potato in the Meghalaya state of India. J Plant Pathol. 2015;97: 135–142. Available: http://www.jstor.org/stable/24579140

179. Gutarra L, Kreuze J, Lindqvist-Kreuze H, De Mendiburu F. Variation of resistance to different strains of *Ralstonia solanacearum* in highland tropics adapted potato genotypes. Am J Potato Res. 2015;92: 258–265. doi:10.1007/s12230-014-9426-4

180. Lopes CA, Rossato M, Boiteux LS. The host status of coffee *(Coffea arabica)* to *Ralstonia solanacearum* phylotype I isolates. Trop Plant Pathol. 2015;40: 1–4. doi:10.1007/s40858-014-0001-9

181. Sikirou R, Zocli B, Paret ML, Deberdt P, Coranson-Beaudu R, Huat J, et al. First report of bacterial wilt of gboma *(Solanum macrocarpon)* caused by *Ralstonia solanacearum* in Benin. Plant Dis. 2015;99: 1640–1640. doi:10.1094/PDIS-02-15-0213-PDN

182. Stulberg MJ, Shao J, Huang Q. A multiplex PCR assay to detect and differentiate Select Agent strains of *Ralstonia solanacearum*. Plant Dis. 2015;99: 333–341. doi:10.1094/PDIS-05-14-0483-RE

183. Stulberg MJ, Huang Q. A TaqMan-based multiplex qPCR assay and DNA extraction method for phylotype IIB sequevars 1 & 2 (Select Agent) strains of *Ralstonia solanacearum*. PLoS One. 2015;10: e0139637. doi:10.1371/journal.pone.0139637

184. Wang H, Huang Y, Wang J, Wang M, Xia H, Lu H. Phenotypic fingerprints of *Ralstonia solanacearum* biovar 3 strains from tobacco and tomato in China assessed by phenotype MicroArray analysis. Plant Pathol J. 2015;14: 38–43. doi:10.3923/ppj.2015.38.43

185. Bhunchoth A, Phironrit N, Leksomboon C, Chatchawankanphanich O, Kotera S, Narulita E, et al. Isolation of *Ralstonia solanacearum*-infecting bacteriophages from tomato fields in Chiang Mai, Thailand, and their experimental use as biocontrol agents. J Appl Microbiol. 2015;118: 1023–1033. doi:10.1111/jam.12763

186. Popoola AR, Ganiyu SA, Enikuomehin OA, Bodunde JG, Adedibu OB, Durosomo HA, et al. Isolation and characterization of *Ralstonia solanacearum* causing bacterial wilt of tomato in Nigeria. Nigerian J Biotechnol. 2015;29: 1–10. doi:10.4314/njb.v29i1.1

187. Lin C-H, Chuang M-H, Wang J-F. First report of Bacterial Wilt caused by *Ralstonia solanacearum* on chard in Taiwan. Plant Dis. 2015;99: 282. doi:10.1094/PDIS-07-14-0715-PDN

188. Shahbaz MU, Mukhtar T, ul-Haque MI, Begum N. Biochemical and Serological Characterization of *Ralstonia solanacearum* Associated with Chilli Seeds from Pakistan. Int J Agric Biol. 2015;17: 31–40. Available: https://www.cabidigitallibrary.org/doi/pdf/10.5555/20153090686

189. Li Y, Feng J, Liu H, Wang L, Hsiang T, Li X, et al. Genetic diversity and pathogenicity of *Ralstonia solanacearum* causing tobacco bacterial wilt in China. Plant Dis. 2016;100: 1288–1296. doi:10.1094/PDIS-04-15-0384-RE

190. Ravelomanantsoa S, Robène I, Chiroleu F, Guérin F, Poussier S, Pruvost O, et al. A novel multilocus variable number tandem repeat analysis typing scheme for African phylotype III strains of the *Ralstonia solanacearum* species complex. PeerJ. 2016;4: e1949. doi:10.7717/peerj.1949

191. Sánchez JAO. Variabilidad genética de la bacteria *Ralstonia solanacearum* de cepas aisladas de plátano en México. Centro de Investigación Científica de Yucatán. 2016. Available: https://pdfs.semanticscholar.org/2a5f/21145faea3ad1b56ddb52117000a45c50268.pdf

192. Ayana G. Diversity and phylotype analysis of *Ralstonia solanacearum* strains causing tomato and potato bacterial wilt in Ethiopia. Pest Management Journal of Ethiopia. 2016 [cited 24 Mar 2021]. Available: https://www.researchgate.net/profile/Muluken_Goftishu/publication/315537001_Evaluation_of_Neem_Seed_and_Citrus_Peel_Powder_for_the_Management_of_Maize_Weevil_Sitophilus_zeamais_Motsch_Coleoptera_Curculionidae_in_Sorghum/links/59079aa20f7e9bc0d59a7d10/Evaluation-of-Neem-Seed-and-Citrus-Peel-Powder-for-the-Management-of-Maize-Weevil-Sitophilus-zeamais-Motsch-Coleoptera-Curculionidae-in-Sorghum.pdf

193. Ailloud F, Lowe TM, Robène I, Cruveiller S, Allen C, Prior P. *In planta* comparative transcriptomics of host-adapted strains of *Ralstonia solanacearum*. PeerJ. 2016;4: e1549. doi:10.7717/peerj.1549

194. Gopalakrishnan C, Artal RB, Sane A. View of Occurrence of *Ralstonia solanacearum* causing bacterial wilt on Bird of Paradise, *Strelitzia reginae* in India. Indian Phytopathol. 2016;69: 44–46. Available: https://epubs.icar.org.in/ejournal/index.php/IPPJ/article/view/71226/30113

195. Guarischi-Sousa R, Puigvert M, Coll NS, Siri MI, Pianzzola MJ, Valls M, et al. Complete genome sequence of the potato pathogen *Ralstonia solanacearum* UY031. Stand Genomic Sci. 2016;11: 7. doi:10.1186/s40793-016-0131-4

196. Ireland KB, Weir BS, Cother EJ, Phantavong S, Phitsanoukane P, Vongvichid K, et al. First report of *Ralstonia pseudosolanacearum* in the Lao PDR. Australas Plant Dis Notes. 2016;11: 36. doi:10.1007/s13314-016-0224-3

197. Jiang Y, Li B, Liu P, Liao F, Weng Q, Chen Q. First report of bacterial wilt caused by *Ralstonia solanacearum* on fig trees in China. For Pathol. 2016;46: 256–258. doi:10.1111/efp.12267

198. Sakthivel K, Gautam RK, Kumar K, Dam Roy S, Kumar A, Devendrakumar C, et al. Diversity of *Ralstonia solanacearum* strains on the Andaman Islands in India. Plant Dis. 2016;100: 732–738. doi:10.1094/PDIS-03-15-0258-RE

199. Guinard J, Vinatzer BA, Poussier S, Lefeuvre P, Wicker E. Draft Genome Sequences of Nine Strains of *Ralstonia solanacearum* Differing in Virulence to Eggplant *(Solanum melongena)*. Genome Announc. 2016;4: e01415–15. doi:10.1128/genomeA.01415-15

200. Kurabachew H, Ayana G. Bacterial Wilt caused by *Ralstonia solanacearum* in Ethiopia: Statusa And Management Approaches: A Review. Int J Phytopathol. 2016;5: 107–119. doi:10.33687/phytopath.005.03.1829

201. Santiago TR, Lopes CA, Caetano-Anollés G, Mizubuti ESG. Phylotype and sequevar variability of *Ralstonia solanacearum* in Brazil, an ancient centre of diversity of the pathogen. Plant Pathol. 2016;66: 383–392. doi:10.1111/ppa.12586

202. Yahiaoui N, Chéron JJ, Jeetah R, Benimadhu S, Félicité J, Cellier G, et al. First Report of *Ralstonia pseudosolanacearum* Phylotype I Causing Bacterial Wilt on Rodrigues Island, Indian Ocean. Plant Dis. 2016;100: 2522. doi:10.1094/PDIS-06-16-0811-PDN

203. Albuquerque GMR, Souza EB, Silva AMF, Lopes CA, Boiteux LS, Fonseca ME de N. Genome sequence of *Ralstonia pseudosolanacearum* strains with compatible and incompatible interactions with the major tomato resistance source Hawaii 7996. Genome Announc. 2017;5. doi:10.1128/genomeA.00982-17

204. Caruso P, Biosca EG, Bertolini E, Marco-Noales E, Gorris MT, Licciardello C, et al. Genetic diversity reflects geographical origin of *Ralstonia solanacearum* strains isolated from plant and water sources in Spain. Int Microbiol. 2017;20: 155–164. doi:10.2436/20.1501.01.298

205. Du H, Chen B, Zhang X, Zhang F, Miller SA, Rajashekara G, et al. Evaluation of *Ralstonia solanacearum* infection dynamics in resistant and susceptible pepper lines using bioluminescence imaging. Plant Dis. 2017;101: 272–278. doi:10.1094/PDIS-05-16-0714-RE

206. Kyaw HWW, Tsuchiya K, Matsumoto M, Iiyama K, Aye SS, Zaw M, et al. Genetic diversity of *Ralstonia solanacearum* strains causing bacterial wilt of solanaceous crops in Myanmar. J Gen Plant Pathol. 2017;83: 216–225. doi:10.1007/s10327-017-0720-0

207. Liu Y, Wu D, Liu Q, Zhang S, Tang Y, Jiang G, et al. The sequevar distribution of *Ralstonia solanacearum* in tobacco-growing zones of China is structured by elevation. Eur J Plant Pathol. 2017;147: 541–551. doi:10.1007/s10658-016-1023-6

208. Rossato M, Santiago TR, Mizubuti ESG, Lopes CA. Characterization and pathogenicity to geranium of Brazilian strains of *Ralstonia* spp. Trop Plant Pathol. 2017;42: 458–467. doi:10.1007/s40858-017-0177-x

209. She X, Yu L, Lan G, Tang Y, He Z. Identification and genetic characterization of *Ralstonia solanacearum* species complex isolates from *Cucurbita maxima* in China. Frontiers in Plant Science. 2017;8. doi:10.3389/fpls.2017.01794

210. Wang L, Wang B, Zhao G, Cai X, Jabaji S, Seguin P, et al. Genetic and pathogenic diversity of *Ralstonia solanacearum* causing potato brown rot in China. Am J Potato Res. 2017;94: 403–416. doi:10.1007/s12230-017-9576-2

211. Yahiaoui N, Chéron J-J, Ravelomanantsoa S, Hamza AA, Petrousse B, Jeetah R, et al. Genetic diversity of the *Ralstonia solanacearum* species complex in the Southwest Indian Ocean Islands. Front Plant Sci. 2017;8: 2139. doi:10.3389/fpls.2017.02139

212. Gutarra L, Herrera J, Fernandez E, Kreuze J, Lindqvist-Kreuze H. Diversity, pathogenicity, and current occurrence of bacterial wilt bacterium *Ralstonia solanacearum* in Peru. Front Plant Sci. 2017;8: 1221. doi:10.3389/fpls.2017.01221

213. Bocsanczy AM, Huguet-Tapia JC, Norman DJ. Comparative genomics of *Ralstonia solanacearum* identifies candidate genes associated with cool virulence. Front Plant Sci. 2017;8: 1565. doi:10.3389/fpls.2017.01565

214. Chen D, Liu B, Zhu Y, Wang J, Chen Z, Che J, et al. Complete genome sequence of *Ralstonia solanacearum* FJAT-1458, a potential biocontrol agent for tomato wilt. Genome Announc. 2017;5. doi:10.1128/genomeA.00070-17

215. Obrador-Sánchez JA, Tzec-Simá M, Higuera-Ciapara I, Canto-Canché B. Genetic diversity of *Ralstonia solanacearum* strains from Mexico associated with Moko disease. Eur J Plant Pathol. 2017;149: 817–830. doi:10.1007/s10658-017-1228-3

216. Norman DJ, Bocsanczy AM, Harmon P, Harmon CL, Khan A. First report of bacterial wilt disease caused by *Ralstonia solanacearum* on blueberries *(Vaccinium corymbosum)* in Florida. Plant Dis. 2017;102: 438–438. doi:10.1094/PDIS-06-17-0889-PDN

217. Sikirou R, Beed F, Ezin V, Hoteigni J, Miller SA. Distribution, pathological and biochemical characterization of *Ralstonia solanacearum* in Benin. Ann Agric Sci. 2017;62: 83–88. doi:10.1016/j.aoas.2017.05.003

218. Tjou-Tam-Sin NNA, van de Bilt JLJ, Westenberg M, Gorkink-Smits PPMA, Landman NM, Bergsma-Vlami M. Assessing the Pathogenic Ability of *Ralstonia pseudosolanacearum (Ralstonia solanacearum* Phylotype I) from Ornamental *Rosa* spp. Plants. Front Plant Sci. 2017;8: 1895. doi:10.3389/fpls.2017.01895

219. Tjou-Tam-Sin NNA, van de Bilt JLJ, Westenberg M, Bergsma-Vlami M, Korpershoek HJ, Vermunt AMW, et al. First report of bacterial wilt caused by *Ralstonia solanacearum* in ornamental *Rosa* sp. Plant Dis. 2017;101: 378–378. doi:10.1094/PDIS-02-16-0250-PDN

220. Kumar S, Kedarnath, Hamsaveni N, Ramanjini Gowda PH, Rohini IB, Rangaswamy KT, et al. Isolation and Characterization of *Ralstonia solanacearum* Causing Bacterial Wilt of Solanaceae Crops. Int J Curr Microbiol App Sci. 2017;6: 1173–1190. doi:10.20546/ijcmas.2017.605.128

221. Bergsma-Vlami M, van de Bilt JLJ, Tjou-Tam-Sin NNA, Westenberg M, Meekes ETM, Teunissen HAS, et al. Phylogenetic assignment of *Ralstonia pseudosolanacearum (Ralstonia solanacearum* phylotype I) isolated from *Rosa* spp. Plant Dis. 2018;102: 2258–2267. doi:10.1094/PDIS-09-17-1345-RE

222. Cho H, Song E-S, Lee YK, Lee S, Lee S-W, Jo A, et al. Analysis of genetic and pathogenic diversity of *Ralstonia solanacearum* causing potato bacterial wilt in Korea. Plant Pathol J. 2018;34: 23–34. doi:10.5423/PPJ.FT.09.2017.0203

223. Ravelomanantsoa S, Vernière C, Rieux A, Costet L, Chiroleu F, Arribat S, et al. Molecular epidemiology of bacterial wilt in the Madagascar highlands caused by Andean (phylotype IIB-1) and African (phylotype III) brown rot strains of the *Ralstonia solanacearum* species complex. Front Plant Sci. 2018;8: 2258. doi:10.3389/fpls.2017.02258

224. She X, He Z, Li H. Genetic structure and phylogenetic relationships of *Ralstonia solanacearum* strains from diverse origins in Guangdong Province, China. J Phytopathol. 2018;166: 177–186. doi:10.1111/jph.12674

225. Shutt VM, Shin G, van der Waals JE, Goszczynska T, Coutinho TA. Characterization of *Ralstonia* strains infecting tomato plants in South Africa. Crop Prot. 2018;112: 56–62. doi:10.1016/j.cropro.2018.05.013

226. Thano P, Akarapisan A. Phylotype and sequevar of *Ralstonia solanacearum* which causes bacterial wilt in *Curcuma alismatifolia* Gagnep. Lett Appl Microbiol. 2018;66: 384–393. doi:10.1111/lam.12857

227. Zhang YW, Chen Y, Hu CH, Li QQ, Lin W, Yuan GQ. Bacterial wilt caused by *Ralstonia pseudosolanacearum* (*R. solanacearum* phylotype I) on *Luffa cylindrica* in China. J Plant Pathol. 2018;100: 593–593. doi:10.1007/s42161-018-0098-7

228. Chesneau T, Maignien G, Boyer C, Chéron J-J, Roux-Cuvelier M, Vanhuffel L, et al. Sequevar diversity and virulence of *Ralstonia solanacearum* Phylotype I on Mayotte Island (Indian Ocean). Front Plant Sci. 2018;8: 2209. doi:10.3389/fpls.2017.02209

229. Weibel J, Tran TM, Bocsanczy AM, Daughtrey M, Norman DJ, Mejia L, et al. A *Ralstonia solanacearum* strain from Guatemala infects diverse flower crops, including new asymptomatic hosts vinca and sutera, and causes symptoms in geranium, mandevilla vine, and new host African daisy *(Osteospermum ecklonis)*. Plant Health Prog. 2018;17: 114–121. Available: https://apsjournals.apsnet.org/doi/abs/10.1094/PHP-RS-16-0001

230. Chamedjeu RR, Masanga J, Matiru V, Runo S. Isolation and characterization of *Ralstonia solanacearum* strains causing bacterial wilt of potato in Nakuru County of Kenya. AJB. 2018;17: 1455–1465. doi:10.5897/AJB2018.16659

231. Montecillo AD, Raymundo AK, Papa IA, Aquino GMB, Jacildo AJ, Stothard P, et al. Near-complete genome sequence of *Ralstonia solanacearum* T523, a phylotype I tomato phytopathogen isolated from the Philippines. Microbiol Resour Announc. 2018;7. doi:10.1128/MRA.01048-18

232. Safni I, Subandiyah S, Fegan M. Ecology, Epidemiology and Disease Management of *Ralstonia syzygii* in Indonesia. Front Microbiol. 2018;9: 419. doi:10.3389/fmicb.2018.00419

233. Jangir R, Sankhla IS, Agrawal K. Characterization, incidence, transmission and biological control of *Ralstonia solanacearum* associated with soybean [*Glycine max* (L.) Merrill] in Rajasthan, India. Res Crops. 2018;19: 472–479. doi:10.31830/2348-7542.2018.0001.18

234. Li X, Huang X, Chen G, Zou L, Wei L, Hua J. Complete genome sequence of the sesame pathogen *Ralstonia solanacearum* strain SEPPX 05. Genes Genomics. 2018;40: 657–668. doi:10.1007/s13258-018-0667-3

235. Singh D, Sinha S, Chaudhary G, Dk Y. Biological characterization and genetic diversity of Indian strains of *Ralstonia solanacearum* biovars 3 and 4 causing bacterial wilt of tomato. J Plant Pathol Microbiol. 2018;09. doi:10.4172/2157-7471.1000443

236. Uwamahoro F, Berlin A, Bucagu C, Bylund H, Yuen J. Potato bacterial wilt in Rwanda: occurrence, risk factors, farmers’ knowledge and attitudes. Food Secur. 2018;10: 1221–1235. doi:10.1007/s12571-018-0834-z

237. Bocsanczy AM, Espindola AS, Norman DJ. Whole-genome sequences of *Ralstonia solanacearum* strains P816, P822, and P824, emerging pathogens of blueberry in Florida. Microbiol Resour Announc. 2019;8. doi:10.1128/MRA.01316-18

238. Izadiyan M, Taghavi SM. Characterization of Iranian *Ralstonia solanacearum* biovar 2 strains by partial sequencing of *egl, mutS* and *pga* genes. Australas Plant Pathol. 2019;48: 607–615. doi:10.1007/s13313-019-00664-w

239. Jimenez Madrid AM, Doyle VP, Ivey MLL. Characterization of *Ralstonia solanacearum* species complex strains causing bacterial wilt of tomato in Louisiana, USA. Can J Plant Pathol. 2019;41: 329–338. doi:10.1080/07060661.2019.1584588

240. Pardo JM, López-Alvarez D, Ceballos G, Alvarez E, Cuellar WJ. Detection of *Ralstonia solanacearum* phylotype II, race 2 causing Moko disease and validation of genetic resistance observed in the hybrid plantain FHIA-21. Trop Plant Pathol. 2019;44: 371–379. doi:10.1007/s40858-019-00282-3

241. Tan X, Qiu H, Li F, Cheng D, Zheng X, Wang B, et al. Complete genome sequence of sequevar 14M *Ralstonia solanacearum* strain HA4-1 reveals novel type III effectors acquired through horizontal gene transfer. Front Microbiol. 2019;10: 1893. doi:10.3389/fmicb.2019.01893

242. Hemelda NM, Safitri R, Suhandono S. Genetic diversity of *Ralstonia solanacearum*, A phytopathogenic bacterium infecting horticultural plants in Java, Indonesia. Biodiversitas. 2019;20: 364–372. doi:10.13057/biodiv/d200209

243. Cho H, Song E-S, Heu S, Baek J, Lee YK, Lee S, et al. Prediction of host-specific genes by pan-genome analyses of the Korean *Ralstonia solanacearum* species complex. Front Microbiol. 2019;10: 506. doi:10.3389/fmicb.2019.00506

244. Mohamed AA, Behiry SI, Younes HA, Ashmawy NA, Salem MZM, Márquez-Molina O, et al. Antibacterial activity of three essential oils and some monoterpenes against *Ralstonia solanacearum* phylotype II isolated from potato. Microb Pathog. 2019;135: 103604. doi:10.1016/j.micpath.2019.103604

245. Kim YS, Lim SR, Kim JW, Lee HJ, Park DH. First Report of *Ralstonia solanacearum* Phylotype I Causing Bacterial Wilt on *Rosa* spp. in Korea. Plant Dis. 2019;103: Not Available. doi:10.1094/pdis-09-18-1604-pdn

246. Pireh S, Harighi B. Phenotypic and genetic properties of Ralstonia solanacearum strains, the causal agent of bacterial wilt of potato isolated from Kurdistan province. 2019. Available: https://www.cabidigitallibrary.org/doi/pdf/10.5555/20193059198

247. Tam HN, Thanh L, Phap TQ, Tung TT, Danh LT, Thi N, et al. Isolation and Virulent Evaluation of *Ralstonia solanacearum* cause the Bacterial Wilt in Chrysanthemum (Chrysanthemum Sp.) from Mekong Delta and Lam Dong Province. Biological Forum– An International Journal. 2019;11: 101–106. Available: https://d1wqtxts1xzle7.cloudfront.net/60912926/2019_Huynh_et_al_Ralstonia_solanacearum_Mekong_Delta_and_Lam_Dong_Province_TRINH_QUANG_PHAP20191015-25409-8n58m0-libre.pdf?1571190876=&response-content-disposition=inline%3B+filename%3DIsolation_and_Virulent_Evaluation_of_Ral.pdf&Expires=1791423478&Signature=LrVKRfKxZmg2eSBnyE-dFsLMHdwytsZIBSMtklVlLxlI99AjgbG3s54KwIem-g81~sRlMjupgQ0d3NkgnS5UZp7G~kd2YpLz2PmJLvjtPk1pj0Zs9N3M3zSlKIwHBVRFhsAR-IrHkC92A-fXDdjzdKOygc0rd8IB4DxyNt7WYfJIN6WYgm7HJ-DgwE20jfpbX~VAAZOZH4dqhTOJZTS~O4s2VfD0GIIjpyRu6Y3d~u-HRYjSYU2jgOwA4D24sL~0pcPZuMj7-8l6qQiQiRrQHeg5gk7zjUy~FtCAAl4OAerEy-UtgHJjy-JfuaO8xF9PZJ6~CJswOSIlK5SMDj8icw&Key-Pair-Id=APKAJLOHF5GGSLRBV4ZA

248. Mutimawurugo MC, Wagara I, Muhinyuza JB, Ogweno J. Virulence and characterization of isolates of potato bacterial wilt caused by *Ralstonia solanacearum* (Smith) in Rwanda. African Journal of Agricultural Research. 2019;14: 311–320. doi:10.5897/AJAR2018.13686

249. Abdurahman A, Parker ML, Kreuze J, Elphinstone JG, Struik PC, Kigundu A, et al. Molecular epidemiology of *Ralstonia solanacearum* species complex strains causing bacterial wilt of potato in Uganda. Phytopathology. 2019;109: 1922–1931. doi:10.1094/PHYTO-12-18-0476-R

250. Freitas RG, Hermenegildo PS, Guimarães LMS, Zauza EAV, Badel JL, Alfenas AC. Detection and characterization of *Ralstonia pseudosolanacearum* infecting *Eucalyptus* sp. in Brazil. For Pathol. 2020;50: e12593. doi:10.1111/efp.12593

251. Lee I, Kim YS, Kim J-W, Park DH. Genetic and pathogenic characterization of bacterial wilt pathogen, *Ralstonia pseudosolanacearum (Ralstonia solanacearum* phylotype I), on roses in Korea. Plant Pathol J. 2020;36: 440–449. doi:10.5423/PPJ.OA.06.2020.0095

252. Pastou D, Chéron JJ, Cellier G, Guérin F, Poussier S. First report of *Ralstonia pseudosolanacearum* phylotype I causing bacterial wilt in New Caledonia. Plant Dis. 2020;104: 278. doi:10.1094/PDIS-05-19-1068-PDN

253. Paudel S, Dobhal S, Alvarez AM, Arif M. Taxonomy and Phylogenetic Research on *Ralstonia solanacearum* Species Complex: A Complex Pathogen with Extraordinary Economic Consequences. Pathogens. 2020;9. doi:10.3390/pathogens9110886

254. Rasoamanana H, Ravelomanantsoa S, Yahiaoui N, Dianzinga N, Rébert E, Gauche M-M, et al. Contrasting genetic diversity and structure among Malagasy *Ralstonia pseudosolanacearum* phylotype I populations inferred from an optimized Multilocus Variable Number of Tandem Repeat Analysis scheme. PLoS One. 2020;15: e0242846. doi:10.1371/journal.pone.0242846

255. Santiago TR, Lopes CA, Caetano-Anollés G, Mizubuti ESG. Genetic structure of *Ralstonia solanacearum* and *Ralstonia pseudosolanacearum* in Brazil. Plant Dis. 2020;104: 1019–1025. doi:10.1094/PDIS-09-19-1929-RE

256. Sedighian N, Krijger M, Taparia T, Taghavi SM, Wicker E, Van Der Wolf JM, et al. Genome resource of two potato strains of *Ralstonia solanacearum* biovar 2 (phylotype IIB sequevar 1) and biovar 2T (phylotype IIB sequevar 25) Isolated from lowlands in Iran. Mol Plant Microbe Interact. 2020;33: 872–875. Available: https://apsjournals.apsnet.org/doi/abs/10.1094/MPMI-02-20-0026-A

257. Vasconez IN, Besoain X, Vega-Celedón P, Valenzuela M, Seeger M. First report of bacterial wilt caused by *Ralstonia solanacearum* phylotype IIB sequevar 1 affecting tomato in different regions of Chile. Plant Dis. 2020;104: 2023. doi:10.1094/pdis-01-20-0181-pdn

258. Alvarez Romero PII, Grabowski Ocampos CJ, Carpio Coba CF, Toro Álvarez SV, Ferreira AT, Mizubuti ESG. First report of *Ralstonia solanacearum* causing bacterial wilt of eucalyptus in Ecuador. Plant Dis. 2020. doi:10.1094/PDIS-11-19-2516-PDN

259. Bihon W, Chen J-R, Kenyon L. Identification and characterization of *Ralstonia* spp. causing bacterial wilt disease of vegetables in Mali. J Plant Pathol. 2020;102: 1029–1039. doi:10.1007/s42161-020-00631-1

260. He Y, Mo Y, Zheng D, Li Q, Lin W, Yuan G. Different sequevars of *Ralstonia pseudosolanacearum* causing bacterial wilt of *Bidens pilosa* in China. Plant Dis. 2020;104: 2768–2773. doi:10.1094/PDIS-12-19-2738-SC

261. Horita M, Sakai YK. Specific detection and quantification of *Ralstonia pseudosolanacearum* race 4 strains from Zingiberaceae plant cultivation soil by MPN-PCR. J Gen Plant Pathol. 2020;86: 393–400. doi:10.1007/s10327-020-00939-x

262. Kunwar S, Bamazi B, Banito A, Carter M, Weinstein S, Steidl OR, et al. First report of bacterial wilt disease of tomato, pepper and gboma caused by the *Ralstonia solanacearum* species complex in Togo. Plant Disease. 2020. doi:10.1094/PDIS-08-20-1665-PDN

263. Mollae A, Hosseinipour A, Azadvar M, Massumi H, Ebrahimi F. Phylotype and sequevar determination and AFLP fingerprinting of *Ralstonia solanacearum* strains causing bacterial wilt of potato in southeastern Iran. Eur J Plant Pathol. 2020;157: 389–402. doi:10.1007/s10658-020-02018-5

264. Patil VU, Vanishree G, Sagar V, Chakrabarti SK. Microsatellites composition in bipartite *Ralstonia solanacearum* genomes: A comparative study between the phylotypes. Indian Phytopathology. 2020;73: 767–770. doi:10.1007/s42360-020-00269-0

265. Sedighian N, Taghavi SM, Hamzehzarghani H, van der Wolf JM, Wicker E, Osdaghi E. Potato-infecting *Ralstonia solanacearum* strains in Iran expand knowledge on the global diversity of brown rot ecotype of the pathogen. Phytopathology. 2020;110: 1647–1656. doi:10.1094/PHYTO-03-20-0072-R

266. da Silva JR, Pais AKL, Albuquerque GMR, Silva AMF, Silva Junior WJ, Balbino V de Q, et al. Genomic sequencing of two isolates of *Ralstonia solanacearum* causing Sergipe facies and comparative analysis with Bugtok disease isolates. Genet Mol Biol. 2020;43: e20200155. doi:10.1590/1678-4685-GMB-2020-0155

267. Prokchorchik M, Pandey A, Moon H, Kim W, Jeon H, Jung G, et al. Host adaptation and microbial competition drive *Ralstonia solanacearum* phylotype I evolution in the Republic of Korea. Microb Genom. 2020;6. doi:10.1099/mgen.0.000461

268. Roman-Reyna V, Truchon A, Sharma P, Hand FP, Mazloom R, Vinatzer BA, et al. Genome resource: *Ralstonia solanacearum* phylotype II sequevar 1 (Race 3 Biovar 2) Strain UW848 from the 2020 U.S. geranium introduction. Plant Dis. 2021;105: 207–208. doi:10.1094/PDIS-06-20-1269-A

269. Sharma K, Kreuze J, Abdurahman A, Parker M, Nduwayezu A, Rukundo P. Molecular diversity and pathogenicity of *Ralstonia solanacearum* species complex associated with bacterial wilt of potato in Rwanda. Plant Dis. 2020;105: 770–779. doi:10.1094/PDIS-04-20-0851-RE

270. Uwamahoro F, Berlin A, Bucagu C, Bylund H, Yuen J. *Ralstonia solanacearum* causing potato bacterial wilt: host range and cultivars’ susceptibility in Rwanda. Plant Pathol. 2020;69: 559–568. doi:10.1111/ppa.13140

271. Gbonamou M, N’Guessan AC, Chen J-R, Kone D, Bihon W, Kenyon L. First phylotype analysis of *Ralstonia solanacearum* causing Eggplant bacterial wilt in the Republic of Guinea. J Anim Plant Sci. 2020;46: 8187–8196. doi:10.35759/janmplsci.v46-2.3

272. Albuquerque GR, Lucena LP, Assunção EF, Mesquita JCP, Silva AMF, Souza EB, et al. Evaluation of tomato rootstocks to *Ralstonia solanacearum* and *R. pseudosolanacearum* in Mata mesoregion, PE. Hortic Bras. 2021;39: 72–78. doi:10.1590/s0102-0536-20210111

273. He Y, Chen Y, Zhang Y, Qin X, Wei X, Zheng D, et al. Genetic diversity of *Ralstonia solanacearum* species complex strains obtained from Guangxi, China and their pathogenicity on plants in the Cucurbitaceae family and other botanical families. Plant Pathol. 2021. doi:10.1111/ppa.13389

274. Siregar BA, Giyanto G, Hidayat SH, Siregar IZ, Tjahjono B. Diversity of *Ralstonia pseudosolanacearum*, the causal agent of bacterial wilt on *Eucalyptus pellita* in Indonesia. Biodiversitas. 2021;22. doi:10.13057/biodiv/d220664

275. Li C, Ju Y, Shen P, Wu X, Cao L, Zhou B, et al. Development of recombinase polymerase amplification combined with lateral flow detection assay for rapid and visual detection of *Ralstonia solanacearum* in tobacco. Plant Dis. 2021;105: 3985–3989. doi:10.1094/PDIS-04-21-0688-RE

276. Pais AKL, Silva JR da, Santos LVSD, Albuquerque GMR, Farias ARG de, Silva Junior WJ, et al. Genomic sequencing of different sequevars of *Ralstonia solanacearum* belonging to the Moko ecotype. Genet Mol Biol. 2021;44: e20200172. doi:10.1590/1678-4685-GMB-2020-0172

277. Freitas RG, Hermenegildo PS, Guimarães LMS, Zauza EAV, Badel JL, Alfenas AC. Molecular characterization and aggressiveness of the *Ralstonia solanacearum* species complex from *Eucalyptus* spp. in Brazil. For Pathol. 2021;51: e12704. doi:10.1111/efp.12704

278. Albuquerque GMR, Silva AMF, Silva JR, Melo EA, Mariano RLR, Lemos MC, et al. Sequevar distribution of *Ralstonia* spp. in Solanaceae in the semiarid climate of the Pernambuco state, Brazil. Eur J Plant Pathol. 2021;159: 13–25. doi:10.1007/s10658-020-02132-4

279. Chen K, Wang L, Chen H, Zhang C, Wang S, Chu P, et al. Complete genome sequence analysis of the peanut pathogen *Ralstonia solanacearum* strain Rs-P.362200. BMC Microbiol. 2021;21: 118. doi:10.1186/s12866-021-02157-7

280. Deberdt P, Cellier G, Coranson-Beaudu R, Delmonteil-Girerd M, Canguio J, Rhino B. First Report of Bacterial Wilt Caused by *Ralstonia solanacearum* on *Plectranthus amboinicus* in Martinique. Plant Dis. 2021;105: 2239. doi:10.1094/PDIS-12-20-2622-PDN

281. Kanyagha H. Characterization of Ralstonia spp. in Tanzania and Potential Integrated Pest Management Strategies for Managing Bacterial Wilt in Tomatoes. Ph.D. Thesis, The Ohio State University. 2021. Available: https://ucdavis.app.box.com/s/1iz648c61dpqaz2l84yhguudrwisbzuk

282. Steidl OR, Truchon AN, Hayes MM, Allen C. Complete genome resources for *Ralstonia* bacterial wilt strains UW763 (phylotype I); Rs5 and UW700 (phylotype II); And UW386, RUN2474, and RUN2279 (phylotype III). Mol Plant Microbe Interact. 2021;34: 1212–1215. doi:10.1094/MPMI-04-21-0086-A

283. Trotereau A, Boyer C, Bornard I, Pécheur MJB, Schouler C, Torres-Barceló C. High genomic diversity of novel phages infecting the plant pathogen *Ralstonia solanacearum*, isolated in Mauritius and Reunion islands. Sci Rep. 2021;11: 5382. doi:10.1038/s41598-021-84305-7

284. Lee SY, Thapa Magar R, Kim HJ, Choi K, Lee S-W. Complete genome sequence of a novel bacteriophage RpY1 infecting *Ralstonia solanacearum* strains. Curr Microbiol. 2021;78: 2044–2050. doi:10.1007/s00284-021-02466-0

285. Akarapisan A, Kumvinit A, Nontaswatsri C, Puangkrit T, Kositratana W. Phylotype, sequevar and pathogenicity of *Ralstonia solanaceaum* species complex from Northern Thailand. J Phytopathol. 2021;170: 176–184. doi:10.1111/jph.13065

286. Bereika MFF, Moharam MHA, Abo-Elyousr KAM, Asran MR. Investigation of virulence diversity in *Ralstonia solanacearum* isolates by a random amplified polymorphic DNA collected from Egyptian potato fields. Archives of Phytopathology and Plant Protection. 2022;55: 1201–1218. doi:10.1080/03235408.2022.2081770

287. Beutler J, Holden S, Georgoulis SJ, Williams D, Norman D, Lowe-Power T. Whole genome sequencing suggests that “non-pathogenicity on banana (NPB)” is the ancestral state of the *Ralstonia solanacearum* IIB-4 lineage. PhytoFrontiers. 2022. doi:10.1094/PHYTOFR-06-22-0068-SC

288. Hossain MM, Masud MM, Hossain MI, Haque MM, Uddin MS, Alam MZ, et al. Rep-PCR analyses reveal genetic variation of *Ralstonia solanacearum* causing wilt of solanaceaous vegetables in Bangladesh. Curr Microbiol. 2022;79: 234. doi:10.1007/s00284-022-02932-3

289. Tan X, Dai X, Chen T, Wu Y, Yang D, Zheng Y, et al. Complete genome sequence analysis of *Ralstonia solanacearum* strain PeaFJ1 provides insights into its strong virulence in peanut plants. Front Microbiol. 2022;13: 830900. doi:10.3389/fmicb.2022.830900

290. Sharma K, Iruegas-Bocardo F, Abdurahman A, Alcalá-Briseño RI, Garrett KA, Goss EM, et al. *Ralstonia* strains from potato-growing regions of Kenya reveal two phylotypes and epidemic clonality of phylotype II sequevar 1 strains. Phytopathology. 2022;112: 2072–2083. doi:10.1094/PHYTO-11-21-0455-R

291. Ghorai AK, Dutta S, Roy Barman A. Genetic diversity of *Ralstonia solanacearum* causing vascular bacterial wilt under different agro-climatic regions of West Bengal, India. PLoS One. 2022;17: e0274780. doi:10.1371/journal.pone.0274780

292. Greenrod STE, Stoycheva M, Elphinstone J, Friman V-P. Global diversity and distribution of prophages are lineage-specific within the *Ralstonia solanacearum* species complex. BMC Genomics. 2022;23: 689. doi:10.1186/s12864-022-08909-7

293. Rincón-Flórez VA, Ray JD, Carvalhais LC, O’Dwyer CA, Subandiyah S, Zulperi D, et al. Diagnostics of banana Blood disease. Plant Dis. 2022;106: 947–959. doi:10.1094/PDIS-07-21-1436-RE

294. Wang K, Chen D, Liu Q, Zhu P, Sun M, Peng D. Isolation and Characterization of Novel Lytic Bacteriophage vB_RsoP_BMB50 infecting *Ralstonia solanacearum*. Curr Microbiol. 2022;79: 245. doi:10.1007/s00284-022-02940-3

295. Soongnern L, Chuapong J, Arunothayanan H, Sratongjun M, Relevante C, de Hoop SJ, et al. Phylotype and sequevar analysis of a *Ralstonia pseudosolanacearum* causing wilt in marigold *(Tagetes erecta)*. J Plant Pathol. 2022;104: 1499–1508. doi:10.1007/s42161-022-01200-4

296. Ariute JC, Felice AG, Soares S, da Gama MAS, de Souza EB, Azevedo V, et al. Characterization and association of Rips repertoire to host range of novel *Ralstonia solanacearum* strains by in silico approaches. Microorganisms. 2023;11: 954. doi:10.3390/microorganisms11040954

297. Aslam MN, Mukhtar T. Distributional spectrum of bacterial wilt of chili incited by *Ralstonia solanacearum* in Pakistan. Bragantia. 2023;82: e20220196. doi:10.1590/1678-4499-2022-0196

298. Baroukh C, Cottret L, Pires E, Peyraud R, Guidot A, Genin S. Insights into the metabolic specificities of pathogenic strains from the *Ralstonia solanacearum* species complex. mSystems. 2023;8: e0008323. doi:10.1128/msystems.00083-23

299. Hasan M, Satoh M, Kiba A, Hikichi Y, Ohnishi K. Complete genome sequences of *Ralstonia solanacearum* strains isolated from infected ginger plants. Microbiol Resour Announc. 2023;12: e0129822. doi:10.1128/mra.01298-22

300. Ding S, Yu L, Lan G, Tang Y, Li Z, He Z, et al. Identification and genomic characterization of *Ralstonia pseudosolanacearum* strains isolated from pepino melon in China. Physiol Mol Plant Pathol. 2023;125: 101977. doi:10.1016/j.pmpp.2023.101977

301. Garcia-Estrada RS, Marquez I, Osuna-Garcia LA, Tovar-Pedraza JM, Cruz-Lachica I. First Report of *Ralstonia pseudosolanacearum* Causing Wilt Disease in Tomato *(Solanum lycopersicum* L.) Plants from Mexico. Plant Dis. 2023. doi:10.1094/PDIS-08-22-1838-PDN

302. Cellier G, Omagwa J, Shem E, Mburu H, Aduwa M, Sekanjako I, et al. First report of *Ralstonia solanacearum* on *Ipomoea hildebrandtii* in Kenya. New Dis Rep. 2023;47: e12163. doi:10.1002/ndr2.12163

303. Le TB, Truong MN, Nguyen BT, Vo DQ, Phan TTP. Draft genome sequencing data of the bacterial wilt, *Ralstonia pseudosolanacearum* T2C-Rasto, from *Cucumis sativus*, in An Giang province, Mekong Delta - Southwest Vietnam. Data Brief. 2023;48: 109252. doi:10.1016/j.dib.2023.109252

304. Nguyen Minh L, Kieu Duc T. Molecular identification of *Ralstonia pseudosolanacearum* causing bacterial wilt in tomatoes from Da Nang by using Colony PCR. CTU J Innov Sustain Dev. 2023;15: 1–7. doi:10.22144/ctu.jen.2023.001

305. Oliver JE, Lewis KJ, Post K, Taylor S, Jogi A. First Report of Bacterial Wilt Disease Caused by *Ralstonia solanacearum* on Southern Highbush Blueberries *(Vaccinium corymbosum* interspecific hybrids) in Georgia U.S.A. Plant Dis. 2022. doi:10.1094/PDIS-07-22-1645-PDN

306. Rasoamanana H, Ravelomanantsoa S, Nomenjanahary M-V, Gauche M-M, Prior P, Guérin F, et al. Bacteriocin production correlates with epidemiological prevalence of phylotype I sequevar 18 *Ralstonia pseudosolanacearum* in Madagascar. Appl Environ Microbiol. 2023;89: e0163222. doi:10.1128/aem.01632-22

307. Valdez-Morales MT, Miranda-Campaña OA, Cruz-Lachica I, Garcia-Estrada RS, Carrillo-Fasio JA, Marquez I, et al. First Report of Bacterial Wilt of Eggplant *(Solanum melongena)* Caused by *Ralstonia pseudosolanacearum* in Mexico. Plant Dis. 2023;107: Not Available. doi:10.1094/PDIS-12-22-2940-PDN

308. Vogelaar M, van de Bilt J, Blom N, Pel C, van Doorn B, Landman M, et al. Presence of *Ralstonia pseudosolanacearum* (phylotype I) in aquatic environments in the Netherlands. Plant Dis. 2023. doi:10.1094/PDIS-11-22-2628-SC

309. Zhao Q, Geng M-Y, Xia C-J, Lei T, Wang J, Cao C-D, et al. Identification, genetic diversity, and pathogenicity of *Ralstonia pseudosolanacearum* causing cigar tobacco bacterial wilt in China. FEMS Microbiol Ecol. 2023;99: fiad018. doi:10.1093/femsec/fiad018

310. Aoun N, Avalos K, Cope-Arguello M, Enriquez C, Nguyen T, Olyushinets A, et al. Genome resource announcement of 3 phylotype I and 14 phylotype II *Ralstonia solanacearum* species complex isolates from South America. Phytofrontiers. 2023;3: 859–862.

311. Traoré O, Wonni I, Cellier G, Boro F, Allibert A, Zombré CT, et al. Genetic and pathogenic diversity of *Ralstonia solanacearum* species complex strains isolated in Burkina Faso. J Phytopathol. 2023;171: 1–11. doi:10.1111/jph.13140

312. Dewberry RJ, Sharma P, Prom JL, Kinscherf NA, Lowe-Power T, Mazloom R, et al. Genotypic and Phenotypic Analyses Show *Ralstonia solanacearum* Cool Virulence is a Quantitative Trait Not Restricted to “Race 3 biovar 2.” Phytopathology. 2024. doi:10.1094/phyto-06-24-0187-r

313. Ding S, Ma Z, Yu L, Lan G, Tang Y, Li Z, et al. Comparative genomics and host range analysis of four *Ralstonia pseudosolanacearum* strains isolated from sunflower reveals genomic and phenotypic differences. BMC Genomics. 2024;25: 191. doi:10.1186/s12864-024-10087-7

314. Galiano-Murillo F, Salas-Lara V, Echandi C, Brenes-Guillén L, Uribe-Lorío L. Draft genomes of two phylotype I and II *Ralstonia solanacearum* species complex (RSSC) isolates causing bacterial wilt in tomato plants from Costa Rica. Microbiol Resour Announc. 2024;13: e0104223. doi:10.1128/mra.01042-23

315. Lin G, Gao J, Zou J, Li D, Cui Y, Liu Y, et al. Whole-genome sequence and characterization of *Ralstonia solanacearum* MLY102 isolated from infected tobacco stalks. Genes (Basel). 2024;15. doi:10.3390/genes15111473

316. Joshi BM, Patel MM, Parmar RB, Patel YR, Singh NK, Tapre PV, et al. Characterization of plant pathogenic *Ralstonia solanacearum* causing bacterial wilt of potato. Plant Arch. 2024;24. doi:10.51470/plantarchives.2024.v24.no.1.048

317. Holeva MC, Glynos PE, Reppa C, Karafla CD. First report of *Ralstonia pseudosolanacearum* causing bacterial wilt on *Rosa* sp. in Greece. Plant Dis. 2024 [cited 7 Oct 2026]. doi:10.1094/PDIS-11-23-2279-PDN

318. Mengting JI, Changjiang C, Liuhe LUO, Zhijian LIN, Menglin Z, Bingye Y, et al. Pathogen identification of kiwi bacterial wilt in Fujian. Journal of Agricultural Science & Technology (1008-0864). 2024;26: 144. Available: https://openurl.ebsco.com/contentitem/gcd:177370035?sid=ebsco:plink:crawler-gcd&id=ebsco:gcd:177370035&crl=c&jrnl=10080864

319. Rana L, Satoh M, Tsuzuki M, Kiba A, Hikichi Y, Ohnishi K. Complete genome sequence of Japanese RSSC phylotype-I strains infecting different host plants. Microbiol Resour Announc. 2024;13: e0048324. doi:10.1128/mra.00483-24

320. Ray JD, Vala B, Mintoff S, Pathania N, Bellgard SE. *Ralstonia solanacearum* species complex in Australia. Plant Dis. 2024;108: 3496–3507. doi:10.1094/PDIS-03-24-0691-SR

321. Rincon-Florez VA, Carvalhais LC, Silva AMF, McTaggart A, Ray JD, O’Dwyer C, et al. Validation of PCR diagnostic assays for detection and identification of all *Ralstonia solanacearum* sequevars causing Moko disease in banana. Phytopathology. 2024 [cited 5 Oct 2024]. doi:10.1094/PHYTO-06-24-0190-R

322. Wang R, Li B, Cai S, Ding Y, Shi M, Jin T, et al. Genetic Diversity of *Ralstonia solanacearum* Causing Tobacco Bacterial Wilt in Fujian Province and Identification of Biocontrol Streptomyces sp. Plant Dis. 2024;108: 1946–1958. doi:10.1094/PDIS-08-23-1604-SR

323. Setiawan AW, Jayanti RM, Herawati MM, Natalia K, Kusuma JGR. Identification and Characterization of *Ralstonia solanacearum* Species Complex from Ginger *(Zingiber* officinale) in Semarang Regency, Indonesia. Agro Bali Agric J. 2024;7: 360–374. doi:10.37637/ab.v7i2.1731

324. Soto-Heredia J, Ramos-Tito S, Alves AR, Ferreira L da C, Calderon LLM, Rossato M. First report of *Ralstonia pseudosolanacearum* causing bacterial wilt in ginger in Peru. Plant Dis. 2024. doi:10.1094/PDIS-09-24-1874-PDN

325. Subedi N, Cowell T, Cope-Arguello M, Paul P, Cellier G, Bkayrat H, et al. Characterization of *Ralstonia pseudosolanacearum* diversity and screening tomato, pepper, and eggplant resistance to manage bacterial wilt in South Asia. PhytoFrontiers^TM^. 2023; PHYTOFR-10-23-0136-R. doi:10.1094/PHYTOFR-10-23-0136-R

326. Wang X, Li C, Huang S, Gao H, Li Y, Chen X, et al. Pathogenic and Comparative Genomic Analysis of *Ralstonia pseudosolanacearum* Isolated from *Casuarina*. Plant Dis. 2024;108: 2809–2819. doi:10.1094/PDIS-01-24-0118-RE

327. Xiao Z, Li G, Yang A, Liu Z, Ren M, Cheng L, et al. Comprehensive genome sequence analysis of *Ralstonia solanacearum* gd-2, a phylotype I sequevar 15 strain collected from a tobacco bacterial phytopathogen. Front Microbiol. 2024;15: 1335081. doi:10.3389/fmicb.2024.1335081

328. Zanon VC, Kafer JM, Hagn J, Cardoso J, Dallemole-Giaretta R, Finatto T, et al. Molecular identification of the causal agent of bacterial wilt in tomato plants in southwestern Paraná. Semin Cienc Agrar. 2024;44: 2249–2258. doi:10.5433/1679-0359.2023v44n6p2249

329. Mountseng CP, Mahot H, Membang G, Mboussi SB, Begoude D, Ambang Z. Morphological and biochemical characterization of strains of *Ralstonia solanacearum*, causal agent of bacterial wilt in tomatoes in Cameroon and screening of virulent strains. Annu Res Rev Biol. 2024;39: 27–42. doi:10.9734/arrb/2024/v39i52079

330. Muradashvili M, Meparishvili G, Sikharulidze Z, Kokashvili T, Ghoghoberidze S, Jabnidze N, et al. PHYLOGENETIC STUDY OF RALSTONIA SOLANACEARUM IN GEORGIA. 2024 [cited 7 Oct 2026]. doi:10.5281/ZENODO.11408419

331. Nakahara H, Mori T, Matsusaki H, Kondo K, Matsuzoe N. Characterization of antibacterial substances produced by phenotypic conversion mutants of *Ralstonia solanacearum* species complex strains of phylotype I and IV. J Plant Pathol. 2024;107: 605–617. doi:10.1007/s42161-024-01810-0

332. Blom NI, Gorkink-Smits PPMA, Landman NM, van de Bilt JLJ, Vogelaar MAW, Raaymakers TM, et al. *Ralstonia solanacearum* (phylotype II) isolated from *Rosa* spp. in the Netherlands is closely related to phylotype II isolates from other sources in the Netherlands and is virulent on potato. Eur J Plant Pathol. 2025;171: 445–458. doi:10.1007/s10658-024-02960-8

333. Gómez AM, Joglekar P, Cline W, Huerta AI. First Report of Bacterial Wilt on Southern Highbush Blueberries *(Vaccinium corymbosum)*, Caused by *Ralstonia solanacearum*, in North Carolina. Plant Dis. 2025;109: 1166. doi:10.1094/PDIS-11-24-2411-PDN

334. Qiu JL, Lin K, Zhang Y, Cai XQ. First Report of *Ralstonia pesudosolanacearum* Causing Bacterial Wilt of *Canna edulis* in China. Plant Dis. 2025 [cited 7 Oct 2026]. doi:10.1094/PDIS-10-24-2174-PDN

335. Bamaniya BS, Sherpa TL, Ashajyothi M, Bhutia LD, Kumar R, Rana M, et al. *Ralstonia solanacearum* isolated from diseased brinjal and tomato: A potential threat to vegetable cultivation in Sikkim, India. Proc Natl Acad Sci India Sect B Biol Sci. 2025;95: 551–557. doi:10.1007/s40011-025-01692-x

336. Balamurugan A, Kumar A, Ashajyothi M, Sakthivel K, Velmurugan S, Charishma K, et al. Phylogenetic diversity and clonal expansion in the *Ralstonia solanacearum* species complex: A case study with bacterial wilt outbreak in eggplant. Plant Pathol. 2025;74: 2963–2979. doi:10.1111/ppa.70068

337. Bergsma-Vlami M, Fornefeld E, Glenz R, Kołodziejska A, Giacobbi F, Gozzi R, et al. Exploring the genetic diversity of recent *Ralstonia pseudosolanacearum* phylotype I findings in Europe. Eur J Plant Pathol. 2026;174: 1121–1135. doi:10.1007/s10658-026-03191-9

338. Chiu M, Lowe-Power T, Grabowski M, Khokhani D. First report of bacterial wilt of ginger caused by *Ralstonia pseudosolanacearum* in the continental United States. Plant Dis. 2025. doi:10.1094/PDIS-11-24-2402-PDN

339. de Oliveira Y de M, Carvalhais LC, Assunção IP, de Andrade Lima GS, dos Santos SL, de Souza EB, et al. *Ralstonia* species infecting Solanaceae and banana occur at different relative abundances in distinct regions of Alagoas state, Brazil. Trop Plant Pathol. 2025;50: 62. doi:10.1007/s40858-025-00749-6

340. Elmgreen VN, Kanyangha H, Cellier G, Bains S, Green S, Hayes L, et al. Genome resource announcement of 19 Tanzanian phylotype I *Ralstonia pseudosolanacearum* draft genomes. PhytoFront. 2025. doi:10.1094/phytofr-11-24-0126-a

341. Horita M, Naito K, Taguchi-Shiobara F, Miyatake K, Sekine D, Kurita K, et al. Complete genome sequences of 96 strains of *Ralstonia solanacearum* species complex phylotype I isolated from solanaceous vegetables in Japan. Microbiol Resour Announc. 2025;14: e0075325. doi:10.1128/mra.00753-25

342. Mogea R, Puspasari A, Ruimassa RMR. Characterization of pathogenic bacteria causing bacterial wilt disease of Kepok banana in Manokwari, west Papua. J Biol Sci Technol Man. 2025;7: 318–325. doi:10.5614/3bio.2025.7.1.4

343. Magar P, Huerta AI, Cellier G, Louws F, Adhikari T. First report of *Ralstonia pseudosolanacearum* phylotype I sequevar 14 causing bacterial wilt on tomato *(Solanum lycopersicum)* and eggplant *(Solanum melongena)* in North Carolina, USA. Plant Dis. 2025;109: 1167. doi:10.1094/PDIS-11-24-2377-PDN

344. Kang J, Sohail MA, Zhang Y, Wang Z, Xu R, Hsiang T, et al. Pan-genome analysis of *Ralstonia pseudosolanacearum* associated with tobacco bacterial wilt in China. BMC Microbiol. 2025;25: 775. doi:10.1186/s12866-025-04485-4

345. Nomenjanahary MV, Rasoamanana H, Javegny S, Rieux A, Ravelomanantsoa S, Poussier S, et al. High-quality genome resource of eight strains of *Ralstonia pseudosolanacearum* causing bacterial wilt in Madagascar. PhytoFront. 2025;5: 94–99. doi:10.1094/phytofr-10-24-0110-a

346. Rezende RR de, Morgan T, Quintero FO, Santos CCD, Oliveira LSS, Mafia RG, et al. First report of *Ralstonia syzygii* causing bacteria wilt on eucalyptus in Brazil. Plant Dis. 2025. doi:10.1094/PDIS-10-24-2229-PDN

347. Saini M, Sagar V, Gupta M, Sharma SK, Saini R. Identification, characterization and genetic diversity of *Ralstonia pseudosolanacearum* causing bacterial wilt of tomato in Himachal Pradesh, India. Physiol Mol Plant Pathol. 2025;138: Not Available. doi:10.1016/j.pmpp.2025.102684

348. Hsu Y-C, Wang C-J, Chung W-H. First report of snow bush wilt caused by *Ralstonia pseudosolanacearum (Ralstonia solanacearum* phylotype I) in Taiwan. Crop Prot. 2025;190: 107070. doi:10.1016/j.cropro.2024.107070

349. Musah SM, Kamiri HW, Birithia RK, Kahariri E. Farmers’ management practices of potato bacterial wilt and its implications in disease prevalence in Kenya. J Agric Ext. 2025;29: 47–62. doi:10.4314/jae.v29i1.6

350. Caly-Simbou E, Nomenjanahary MV, Javegny S, Boyer C, Chéron J-J, Ramin-Mangata S, et al. First genomic analysis of a strain of *Ralstonia pseudosolanacearum* isolated from Mayotte island. BMC Genom Data. 2025;27: 10. doi:10.1186/s12863-025-01395-2

351. Kanyagha H, Miller S, Mbije N. Potential of host resistance as an important tool in the management of bacterial wilt in tomatoes. East African Journal of Science, Technology and Innovation. 2026;7. doi:10.37425/xp6rhn49

352. Pirc M, Radišek S, Pavlič Nikolić E, Dreo T. First Report of *Ralstonia pseudosolanacearum* (Phylotype I) Causing Bacterial Wilt in *Zingiber officinale* in Slovenia. Plant Dis. 2026;110: 1002. doi:10.1094/pdis-10-25-2046-pdn

353. Sikirou R, Dossoumou M-E, Cellier G, Souna DA, De Troij A, Deberdt P. First Report of Bacterial Wilt on African Basil (*Ocimum gratissimum*) Caused by *Ralstonia pseudosolanacearum* Phylotype I in Benin. Plant Dis. 2026;110: 999. doi:10.1094/pdis-10-25-2043-pdn

354. Obregon V, Shin GY, Galdeano E, Escobar R, Lattar T, Ibañez JM, et al. Phylogenomic Taxonomic Analysis of Ralstonia solanacearum Strains causing Bacterial Wilt Disease in Northeastern Argentina. 2026. p. 001307.v1. doi:10.1099/acmi.0.001307.v1

355. Sangsil P, Nualsri C, Insaringkan W, Woraathasin N, Plodpai P, Nakkanong K. Molecular Characterization, Virulence Diversity, and Host Genome-Dependent Resistance to *Ralstonia syzygii* subsp. celebesensis Causing Banana Blood Disease in Southern Thailand. Research Square. Research Square; 2025. doi:10.21203/rs.3.rs-7803689/v1

356. Price MN, Dehal PS, Arkin AP. FastTree 2--approximately maximum-likelihood trees for large alignments. PLoS One. 2010;5: e9490. doi:10.1371/journal.pone.0009490

357. Letunic I, Bork P. Interactive Tree Of Life (iTOL) v5: an online tool for phylogenetic tree display and annotation. Nucleic Acids Res. 2021;49: W293–W296. doi:10.1093/nar/gkab301

